# SNHG10 promote tumorigenesis via miR-150/VEGFA/EGFR/AKT/ERK/mTOR axis and gemcitabine resistance in PDAC

**DOI:** 10.1101/2025.04.24.650424

**Authors:** Gouri Pandya, Aishwarya Singh, Suman Sourav, Sharon Raju, Rachana Kumari, Rashi Sharma, Vidhi Goyal, Bhudev C Das, Gautam Sethi, Amit Kumar Pandey, Deepti Pandita, Rajender K Motiani, Manoj Garg

**Affiliations:** Amity Institute of Molecular Medicine and Stem Cell Research (AIMMSCR), Amity University, Uttar Pradesh, Sector-125, Noida-201313, India; Laboratory of Calciomics and Systemic Pathophysiology (LCSP), Regional Centre for Biotechnology (RCB), Faridabad-121001, India; Department of Histopathology, Pathology and Laboratory Medicine, Medanta Hospital, Gurgaon, India; Amity Foundation for Science, Technology and Innovation Alliances, Amity University, Uttar Pradesh, Sector-125, Noida-201313, India; Department of Pharmacology, Yong Loo Lin School of Medicine, National University of Singapore, Singapore, 117600, Singapore; Department of Biotechnology, National Institute of Pharmaceutical Education & Research (NIPER) Ahmedabad, Gandhinagar-382355, India; Delhi Institute of Pharmaceutical Sciences & Research (DIPSAR), Delhi Pharmaceutical Sciences & Research University (DPSRU), New Delhi-110017, India

**Keywords:** Pancreatic ductal adenocarcinoma, small nucleolar RNA host gene 10, VEGF-A, EGFR, AKT, ERK1/2, miR-150-5p, gemcitabine resistance, cell cycle, apoptosis, and xenograft model

## Abstract

SNHG10 emerged as a key regulator in progression and metastasis of cancers. However, potential of SNHG10 in PDAC tumorigenesis, gemcitabine resistance, and underlying mechanisms remains poorly understood. We observed significant upregulation of SNHG10 in 179 PDAC cases, revealing a positive correlation with clinical stages. Our results showed a significant SNHG10 overexpression in several PDAC cells. Downregulation of SNHG10 significantly decreased the proliferation, clonogenicity, EMT, tumor growth in the xenograft model, and the induction of cell cycle arrest and apoptosis of PDAC cells. Mechanistically, SNHG10 knockdown significantly inhibited the expression of vimentin, N-cadherin, survivin, CDK4, CDK6, cyclin B1, cyclin D1, aurora kinase A, and B, with an increased expression of E-cadherin and p21. RNA Immunoprecipitation data displayed physical interaction among SNHG10, miR-150-5p, and VEGFA in PDAC cells. SNHG10 silencing led to the significant induction of miR-150-5p, which repressed VEGFA expression in PDAC cells. SNHG10 downregulation enhanced gemcitabine sensitivity in PDAC cells. SNHG10 silencing suppressed the phosphorylation of EGFR, AKT, ERK1/2, mTOR, and c-MET pathways. Silencing SNHG10 and its regulated signalling offers a novel prospective therapeutic strategy.

## Introduction

Pancreatic ductal adenocarcinoma (PDAC) is one of the most lethal cancers because of its late diagnosis, highly metastatic, drug-resistant, and aggressive nature(Cortesi et al, 2024; Sha et al, 2024). The 5-year survival of pancreatic cancer patients is less than 8%(Kirtonia et al, 2022; Pandya et al, 2020; Sha et al, 2024; Zhou et al, 2024). The surgical resection followed by chemotherapy is the first choice of treatment for PDAC cases in the early stages and is often associated with a positive long-term outcome. Nab-paclitaxel plus gemcitabine, 5-fluorouracil, and FOLFIRINOX are FDA-approved for treating advanced pancreatic carcinoma(Chien et al, 2014; Chien et al, 2018; Pandya et al, 2020; Pandya et al, 2022). However, the patients with advanced pancreatic carcinoma displayed a high rate of inherent resistance to these chemotherapeutic drugs, which posed a great challenge in the clinics. Currently, robust biomarkers for early diagnosis, prognosis, and therapy against PDAC are lacking. Thus, the identification and development of highly sensitive, specific, and reliable biomarkers for diagnostic and prognostic benefits, along with effective molecular therapeutic targets, are urgent needs for improving the survival outcome and management of PDAC patients. Over the last decade, advancements in next-generation sequencing technologies have provided more depth and knowledge to understand human genomes(Lander et al, 2001). Recently, transcriptomics studies using human tissue samples have identified a variety of functional non-coding RNA, including microRNA (miRNA), long non-coding RNA (lncRNA), small interfering RNA, small nucleolar RNA, and Piwi-interacting RNA(Palazzo & Koonin, 2020).

The lncRNAs are > 200 nucleotide nonprotein coding sequences, structurally showing a close resemblance with the mRNA transcription start site, polyadenylated tails at 3′, capping at 5′, and active splicing machinery to produce a final product(Kanojia et al, 2022; Shabna et al, 2023; Yadav et al, 2021). The small nucleolar RNA host gene (SNHG) belongs to the family of lncRNA(Huang et al, 2022). Emerging studies have shown that SNHG(s) can regulate important cellular functions to maintain cellular homeostasis. However, the dysregulated expression of SNHG(s) has been observed in several diseases, including diabetes, neurological disorders, and human malignancies(Xiao et al, 2023). SNHGs were shown to be involved in cell proliferation, cell cycle progression, tumor immune environment, invasion, and metastasis of cancer cells, acting as an oncogene or a tumor suppressor depending on the origin of cancers(Williams & Farzaneh, 2012; Xiao et al, 2023). Specifically, the SNHG10 is well characterized and displayed to be overexpressed in various cancers, including acute myeloid leukaemia(Xiao et al, 2021), gastric cancer(Yuan et al, 2021), glioma(Jin et al, 2020), colorectal cancer(Zhang et al, 2021), osteosarcoma(He et al, 2020; Zhu et al, 2020), and prostate cancer(Cao et al, 2021). The overexpression of SNHG10 was found to be associated with inferior overall survival in acute myeloid leukemia(Xiao et al, 2021), hepatocellular carcinoma(Lan et al, 2019), and colorectal cancer(Zhang et al, 2021). On the contrary, several functional studies have confirmed the downregulation of SNHG10 and its tumor suppressive function in lung adenocarcinoma(Liang et al, 2020; Zhang et al, 2020) and ovarian cancer(Lv et al, 2022). Single-cell RNA sequencing data have displayed that SNHG10, SPP1, CASC19, LINC00683, and LINC00237 are involved in immune signatures for developing a prognostic model(Chen et al, 2022b). Several studies have revealed that SNHG10 controls tumorigenesis by altering molecular mechanisms, including epigenetic regulation, sponging of miRNA, and signaling pathways(Zhu et al, 2025).

Henceforth, there is a lack of understanding regarding the potential role and molecular mechanism of SNHG10 in the tumorigenesis and chemotherapeutic resistance in PDAC. In this study, we reported the robust expression of the SNHG10 transcript in PDAC tissues and cell lines. The downregulation of SNHG10 displayed reduced cell viability, proliferation, clonogenicity, migration, EMT, and tumor growth in the PDAC xenograft. Silencing of SNHG10 sensitized the gemcitabine-resistant PDAC cells. Collectively, this study uncovered that SNHG10 modulates the miR-150-5p/VEGF-A/EGFR/AKT/ERK1/2/mTOR signalling cascade, which may be helpful for the development of new strategies for PDAC treatment.

## Results

### SNHG10 is highly expressed in PDAC patient specimens that correlated with clinical stages and in a panel of PDAC cell lines

We analyzed the expression of the SNHG10 in the PDAC and normal pancreatic transcriptomics datasets available in TCGA and The Genotype-Tissue Expression [GTEx; https://www.gtexportal.org] containing 179 PDAC and 171 normal pancreatic samples using the UALCAN [http://ualcan.path.uab.edu/], and GEPIA, Gene Expression Profiling Interactive Analysis [http://gepia.cancer-pku.cn/]. The analysis revealed that the SNHG10 transcripts were highly expressed in 179 pancreatic cancer patients than in 171 normal pancreatic tissues **(Fig. 1a)**. The elevated level of SNHG10 was positively associated with the pathological stages of the PDAC **(Fig. 1b)**. Furthermore, we validated the expression of SNHG10 transcripts in a panel of seven PDAC and hTERT-HPNE (human normal pancreatic epithelial) cells. The qRT-PCR results showed significantly higher expression of SNHG10 in SW1990, PANC-1, Panc10.05, AsPC-1, BxPC-3, and CFPAC-1 cells as compared to that in hTERT-HPNE cells **(Fig. 1c)**. No significant change in the SNHG10 expression was observed in MIA PaCa-2 cells **(Fig. 1c)**.

**Fig. 1.**
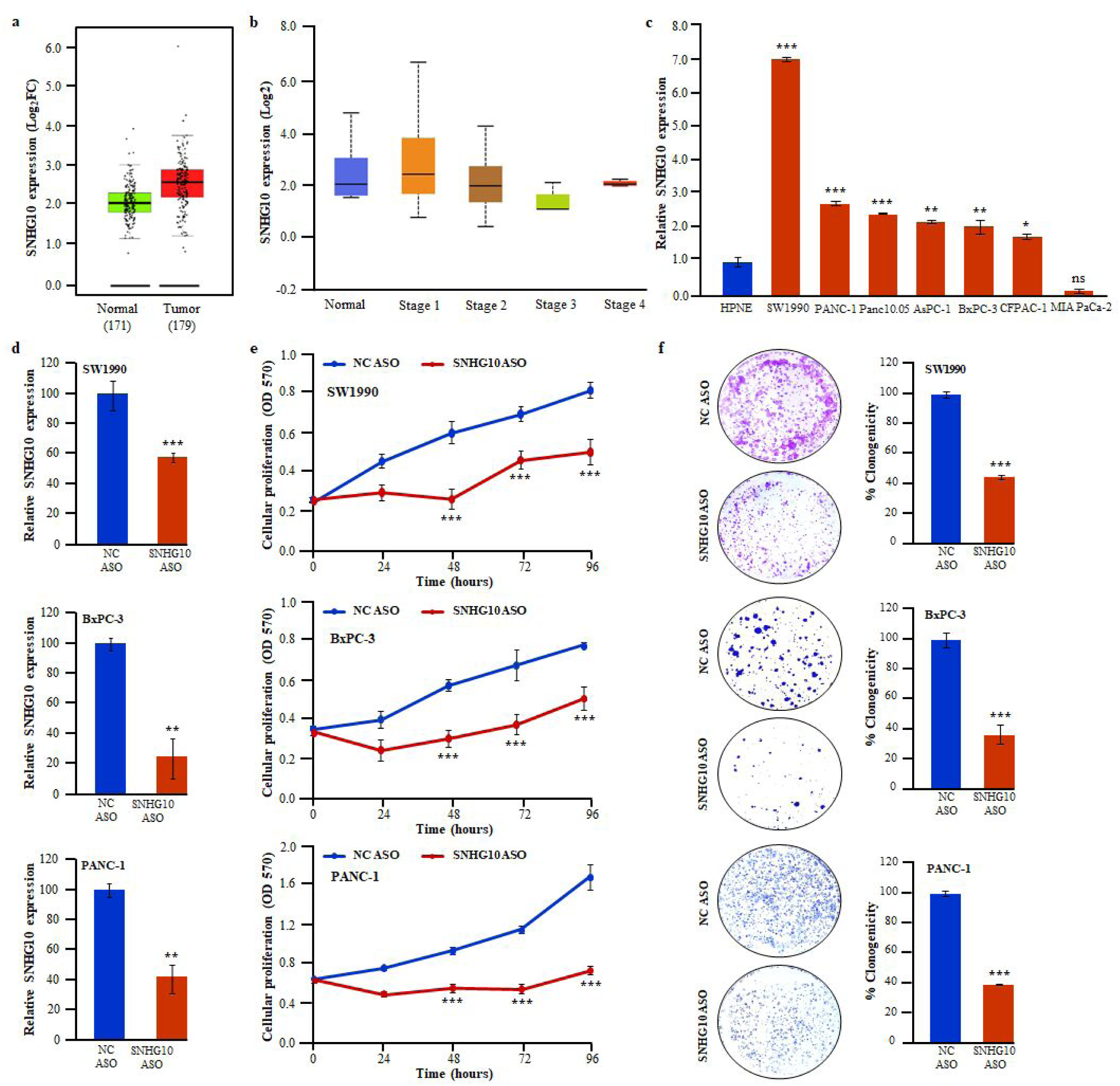
SNHG10 is highly expressed in PDAC patient specimens, in a panel of PDAC cell lines, and downregulation of SNHG10 significantly suppresses the viability and clonogenicity of PDAC cells in vitro. **(a)** Transcriptomic analysis of the TCGA 179 PDAC and 171 normal pancreatic RNA sequencing datasets for the expression of SNHG10 using GEPIA. **(b)** Overexpression of SNHG10 transcript in clinical stages of PDAC. **(c)** The qRT-PCR data revealed higher expression of SNHG10 transcript in PDAC cells than in hTERT-HPNE cells. **(d)** Transfection of SNHG10 ASO and qRT-PCR was used for knockdown of SNHG10 in SW1990, BxPC-3, and PANC-1 cell lines. **(e)** SNHG10 downregulation caused a significant reduction in the cell proliferation ability of SW1990, BxPC-3, and PANC-1 cells as determined by MTT assay. **(f)** Silencing of SNHG10 reduces the clonogenic ability of PDAC cells. All the experiments were performed three times in biological triplicate. The results are presented as the means ± SDs; n=3. *p < 0.05, **p < 0.01, ***p < 0.001; two-tailed Student t test.

### Downregulation of SNHG10 expression significantly suppressed the viability and clonogenicity of PDAC cells

Antisense nucleotides (ASOs) are emerging as an important tool for the effective knockdown and therapeutic targeting of lncRNA. The lncRNA-specific ASOs induce the degradation of targeted lncRNA transcripts by triggering RNase-H-mediated cleavage(Lee & Mendell, 2020). To achieve a successful knockdown of SNHG10 transcript expression, the SW1990, BxPC-3, and PANC-1 cells were transfected with SNHG10-specific ASOs. The qRT-PCR analysis confirmed that SNHG10 ASO treatment caused significant reduction in SNHG10 expression compared to negative control (NC) ASO in SW1990, BxPC-3, and PANC-1 cells **(Fig. 1d)**. The downregulation of the SNHG10 transcript significantly decreased the cellular proliferation of SW1990, BxPC-3, and PANC-1, cells **(Fig. 1e)**.

Furthermore, the downregulation of SNHG10 displayed a significant decrease in clonogenic growth compared to NC ASO-treated SW1990, BxPC-3, and PANC-1 cells. Notably, the size and the number of colonies were markedly decreased in the SNHG10 ASO group **(Fig. 1f)**. Additionally, we have used the siRNA approach to silence the expression of the SNHG10 transcript to reconfirm the phenotypic effect on the viability and clonogenicity as was observed upon ASO-mediated downregulation of SNHG10. The results of siRNA-mediated knockdown of SNHG10 lead to the significant suppression of viability **(Supplementary Fig. S1a)** and clonogenicity **(Supplementary Fig. S1b)** of SW1990, PANC-1, and BxPC-3 cells compared to the negative target (NT) siRNA. Importantly, there is no significant difference in the proliferation **(Supplementary Fig. S2a)** and clonogenic ability **(Supplementary Fig. S2b)** of the hTERT-HPNE cells transfected with SNHG10 ASO, indicating that SNHG10 ASO does not affect the phenotype of the human normal pancreatic cells whereas SNHG10 downregulation has marked effect on the viability of pancreatic cancer cells without producing any off-target effect. Collectively, findings suggested that SNHG10 acts as an oncogenic lncRNA in the PDAC.

### Downregulation of SNHG10 inhibited cell migration via repression of epithelial-to-mesenchymal transition in PDAC cells

Migration of the cancerous cell from the original tumor site is responsible for metastasis, drug resistance, and recurrence(Fares et al, 2020; Lambert et al, 2017). The downregulation of SNHG10 significantly inhibited the migration of SW1990, PANC-1, and BxPC-3 cells through the Boyden chamber compared with NC ASO or NT siRNA **(Fig. 2a, b)**. Further, we evaluated the effect of SNHG10 depletion on the molecular mechanism associated with epithelial-to-mesenchymal transition (EMT). Mechanistically, our data revealed that depletion of SNHG10 led to a significant increase in the E-cadherin protein expression, a critical biomarker for maintaining epithelial phenotype, as well as the reduction in protein expression of vimentin and N-cadherin (mesenchymal markers) in SW1990 and PANC-1 **(Figure 2c, d)**. Noticeably, the downregulation of SNHG10 has no significant effect on the migratory ability of the hTERT-HPNE cells **(Supplementary Fig. S2c)**. Thus, our data indicated that the downregulation of SNHG10 lncRNA suppressed the migration of PDAC cells through the reversal of EMT.

**Fig. 2.**
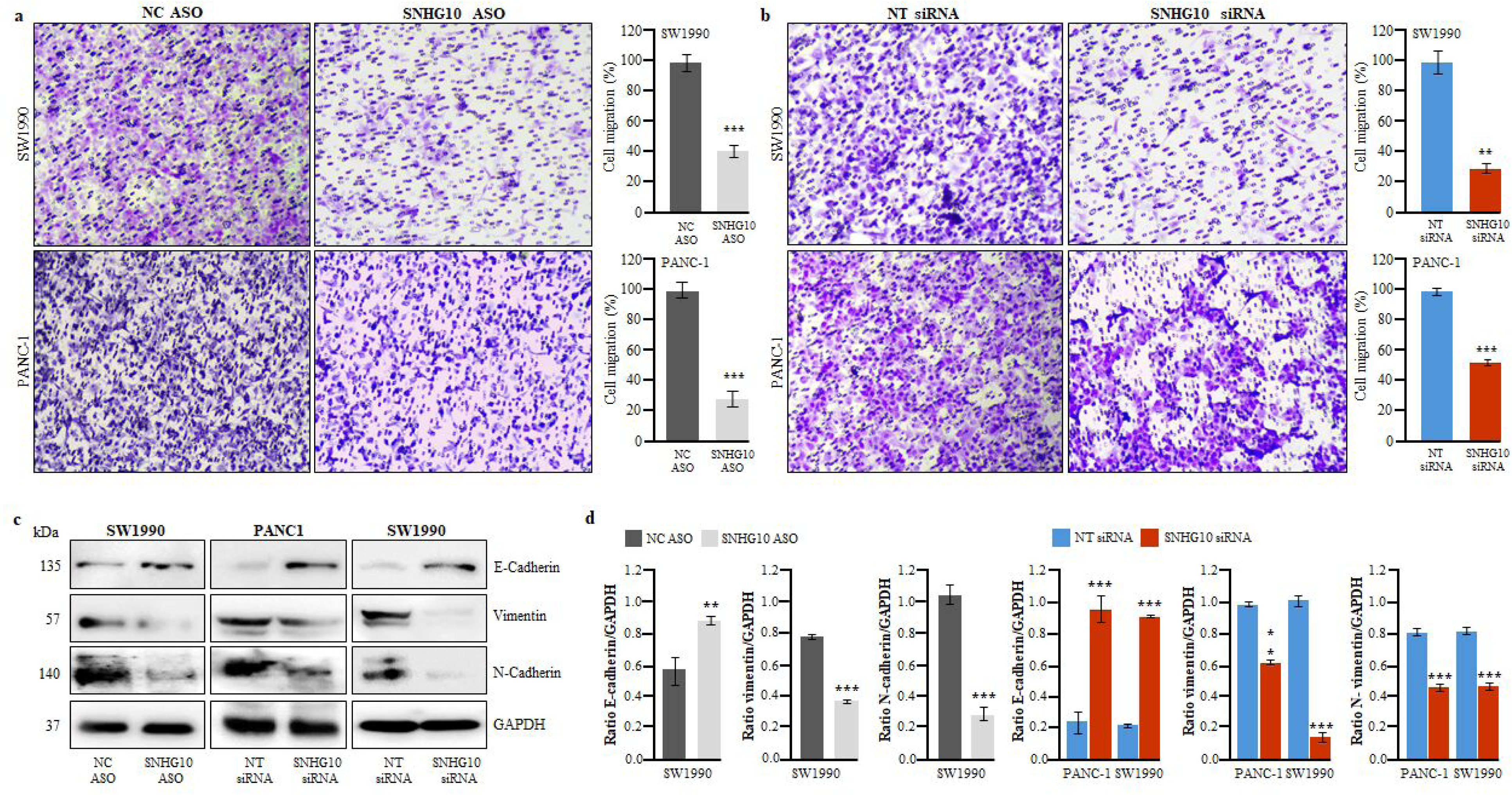
Downregulation of SNHG10 inhibited cell migration via repression of epithelial-to-mesenchymal transition in PDAC cells. **(a** and **b)** Boyden chamber assay determined the effect of SNHG10 downregulation on the migration ability of SW1990 and PANC-1 cells. (**c)** Western blotting experiments displayed the expression of E-cadherin, vimentin, and N-cadherin in SNHG10-depleted SW1990 and PANC-1 cells. **(d)** Densitometry analysis of western blotting data by ImageJ software. Student t-test was performed for statistical significance as shown by means ± SDs; n=3, *p < 0.05, **p < 0.01, *** p < 0.001.

### Downregulation of SNHG10 induced cell cycle arrest and apoptosis in PDAC cells

Next, to investigate the involvement of SNHG10 lncRNA in the induction of cell cycle arrest and apoptosis, flow cytometry analysis was performed. The cell cycle analysis results have displayed that the downregulation of SNHG10 significantly arrested the SW1990, PANC-1, and BxPC-3 cells in the G2/M phase and subG1 phase leading to a decrease in the cell population during the G0/G1 phase of the cell cycle **(Fig. 3a, b; Supplementary Fig. 3a)**. Furthermore, our data demonstrated that the downregulation of SNHG10 resulted in significantly increased early and late apoptosis in SW1990, PANC-1, and BxPC-3 cell lines **(Fig. 3c, d; Supplementary Fig. 3b)**. To explore the mechanism(s) underlying growth inhibition, cell cycle arrest, and apoptosis, we performed protein expression analysis of cell cycle and apoptosis regulators and discovered that the depletion of SNHG10 expression significantly repressed the protein level of cyclin D1, and associated cyclin-dependent kinase-4 and −6 (CDK4, and CDK6) leading to disruption of cyclin D1-CDK4/CDK6 complex in SW1990, and PANC-1 cells **(Fig. 3e)**. Additionally, the silencing of SNHG10 significantly restored the expression of tumor suppressor protein p21 along with the suppressed expression of cyclin B1, survivin, aurora kinase A, and aurora kinase B in SW1990 and PANC-1 cells **(Fig. 3e; Supplementary Fig. S4)**. These findings together indicated that SNHG10 lncRNA is a critical regulator in the molecular processes involved in the cell cycle and survival of PDAC cells.

**Fig. 3.**
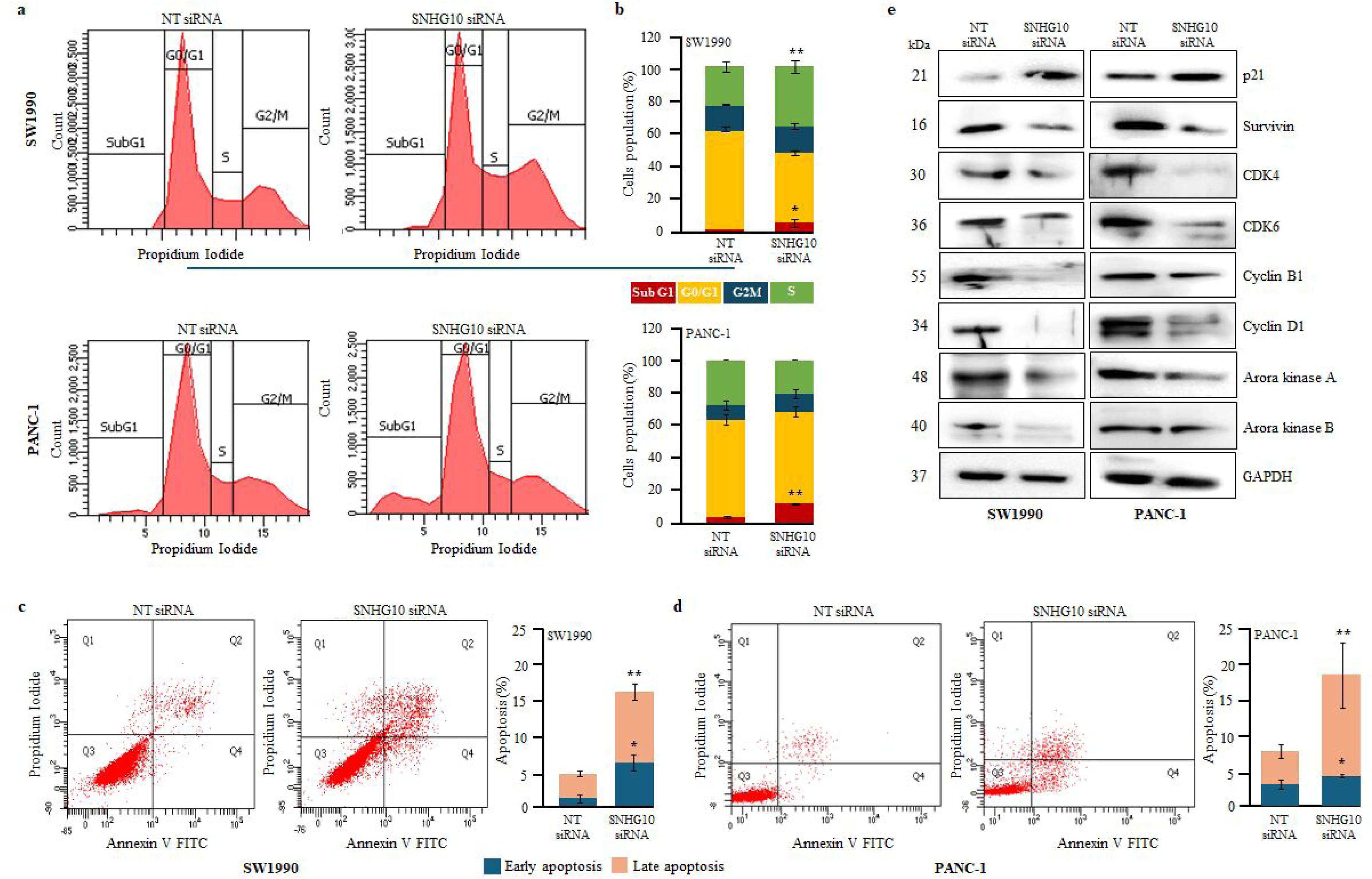
Downregulation of SNHG10 induced cell cycle arrest and apoptosis in PDAC cells. **(a)** Cell cycle distribution measured by propidium iodide staining in SW1990 and PANC-1 cells transfected with SNHG10 siRNA and NT-siRNA. (**b)** Quantification and histogram represented the different phases of the cell cycle. (**c** and **d)** Apoptosis was determined using Annexin V and propidium iodide staining in SW1990 and PANC1 cells transfected with SNHG10 siRNA and NT siRNA. **(e**) Cell lysates were prepared for Western blot analysis of the regulator proteins involved in cell cycle and apoptosis. GAPDH was used as a loading control. All the experiments were performed three times in biological triplicate. The results are presented as the means ± SDs; n=3. *p < 0.05, **p < 0.01, ***p < 0.001; two-tailed Student t test.

### SNHG10 interacted with miR-150-5p and modulated the expression of VEGF-A in PDAC cells

Emerging pieces of evidence indicate that lncRNA can serve as competing endogenous RNA to sequester the expression of miRNAs. This sponging of miRNAs has been observed in modifying the mRNA expression of miRNA-targeted genes(Pandya et al, 2020). Therefore, the bioinformatics analysis was carried out using several tools, including miRcode, lncRNASNP2, LncBase, and Starbase to find out the potential miRNA targets of SNHG10 lncRNA. The Lncbase and miRcode tools predicted the high binding affinity among miR-150-5p and SNHG10 **(Fig. 4a).** The miR-150-5p displayed 7 to 8 Mer seed sequences conservation with SNHG10 **(Fig. 4a).** Further, we analysed the expression of miR-150-5p in pancreatic cancer tissues and the data revealed the marked downregulation of miR-150-5p in patients with PDAC compared to normal pancreatic tissues **(Fig. 4b)**. The downregulation of miR150-5p was consistent in both the stages and grades in PDAC patients **(Fig. 4c and d).** Additionally, the qPCR data displayed significant downregulation of miR-150-5p in SW1990 and BxPC-3 cells compared to hTERT-HPNE cells **(Fig. 4e)**, suggesting that miR-150-5p can act as tumor suppressor in PDAC. Interestingly, the silencing of SNHG10 significantly restored the expression of miR-150-5p in both SW1990 and BxPC-3 cells, indicating that SNHG10 could potentially sponge miR-150-5p **(Fig. 4f)**. This is a well-known fact that miR can efficiently target protein-coding genes to control the posttranscriptional regulation to maintain cellular equilibrium. Henceforth, mirTarbase, Starbase, and miRDB were used to search for a potential target of miR150-5p. A total of 16 potential protein-coding target genes were commonly predicted across three databases **(Fig. 4g).** Among them, VEGF-A was selected as a target of miR150-5p due to its importance and association with tumorigenesis in several human cancers. We noticed that VEGF-A is robustly expressed in patients with PDAC **(Fig. 4h).** Previously, studies have also observed the interaction of miR-150-5p with VEGF-A in colon tumors(Chen et al, 2021; Chen et al, 2018). As expected, the coexpression analysis showed a significant negative correlation between miR-150-5p and VEGF-A in PDAC tumor tissues **(Fig. 4i)**. Next, we were curious to know the correlation between SNHG10 and VEGF-A. Interestingly, our data analysis showed a significant positive correlation between SNHG10 and VEGF-A in PDAC **(Fig. 4j)**. Furthermore, our data confirmed that silencing of SNHG10 significantly decreases the expression of VEGF-A both at transcription (mRNA) and translational (protein) level in SW1990 and BxPC-3 cells **(Fig. 4k and 4l).** Next, we checked whether the interaction between SNHG10 and miR-150-5p is critical to regulate the VEGF-A expression using Ago2 (the core component of the RISC) RNA immunoprecipitation assays. The results of the RIP assay revealed that SNHG10, miR-150-5p, and VEGF-A are significantly enriched and interacted in the Ago2 RIP group compared with IgG control in SW1990 and BxPC-3 cells **(Fig. 4m and 4n)**. Taken together, these data suggested that SNHG10, miR-150-5p, and VEGF-A physically existed in the RISC complex, indicating that silencing SNHG10 could upregulate tumor suppressive miR-150-5p, which in turn repressed VEGF-A through sponging.

**Fig. 4.**
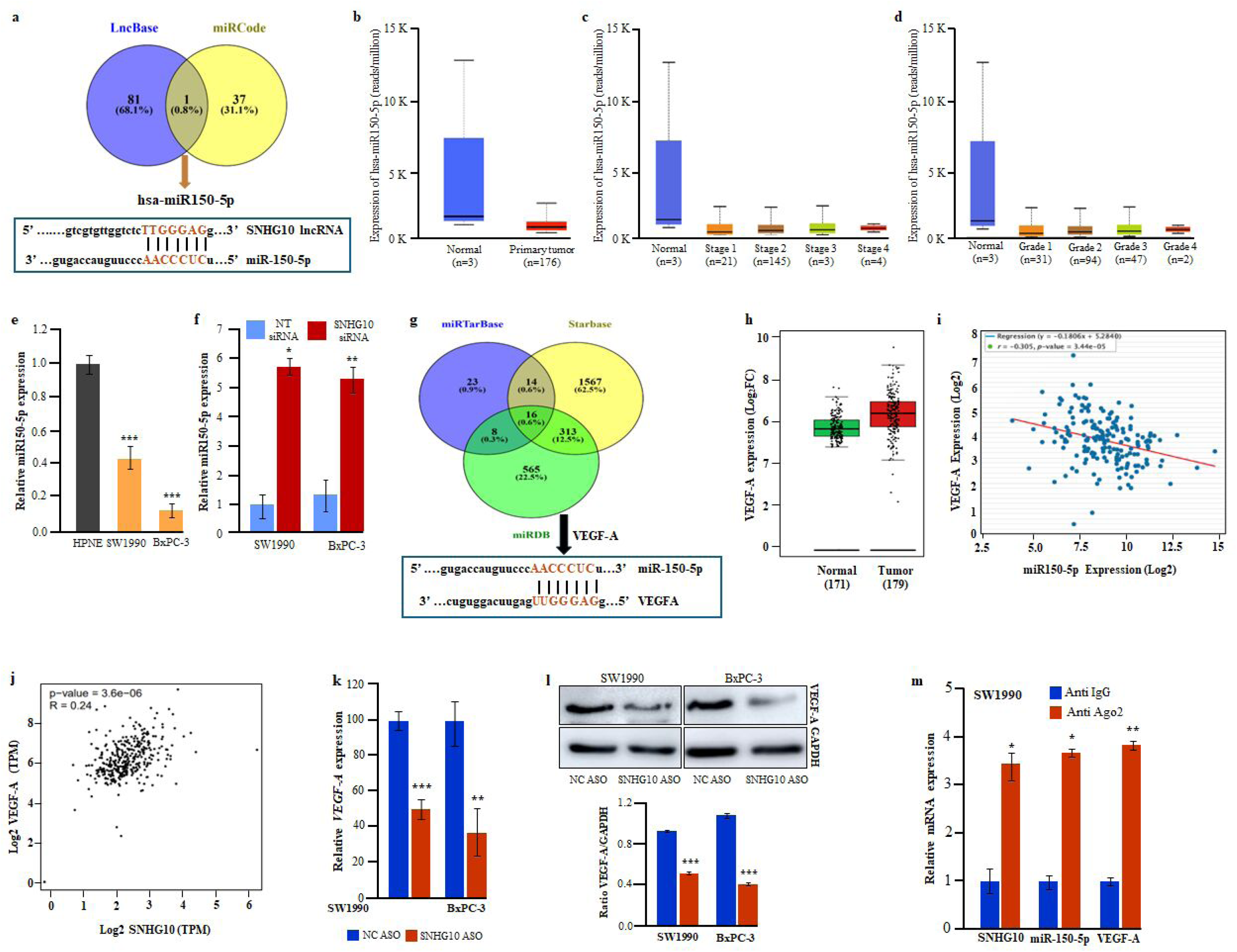
SNHG10 interacted with miR-150-5p and modulated the expression of VEGF-A in PDAC cells. **(a)** The Venn diagram displayed miR-150-5p as a target of SNHG10 predicted by Lncbase and miRCode, and binding sites between SNHG10 and miR-150-5p predicted using bioinformatics analysis. **(b** to **d)** The expression of miR-150-5p and its correlation with pathological stages and grades in PDAC TCGA data. **(e)** The qRT-PCR analysis revealed downregulation of miR-150-5p in SW1990 and BxPC-3 cells. **(f)** Knockdown of SNHG10 restored the expression of miR-150-5p in SW1990 and BxPC-3 cells, as shown by qRT-PCR. **(g)** The miRTarbase, starbase, and miRDB databases predicted VEGF-A as the target genes of miR-50-5p, as shown by the Venn diagram, and conservation in binding among miR-150-5p and VEGF-A. **(h)** TCGA data analysis confirmed overexpression of VEGF-A in PDAC. **(i)** Negative correlation was noticed between miR-150-5p and VEGF-A using StarBase. **(j)** The scattered plot showed **the** positive correlation between SNHG10 and VEGF-A using GEPIA analysis of the PDAC TCGA data. **(k** and **l)** qRT-PCR data demonstrated the downregulation of SNHG10 repressed VEGF-A expression at the mRNA and protein level in SW1990 and BxPC-3 cells. The densitometry analysis displayed significant inhibition of VEGF-A. **(m)** RNA immunoprecipitation using anti-Ago2 detected the interaction of SNHG10/miR-150-5p/VEGF-A in SW1990 cells. The results are presented as the means ± SDs; n=3. *p < 0.05, **p < 0.01, ***p < 0.001; two-tailed Student t test.

### Downregulation of SNHG10 suppressed EGFR/AKT/ERK1/2/mTOR signalling cascade in PANC-1 and SW1990 cells

EGFR, epidermal growth factor receptor signalling, plays an important role in tumor growth, survival, metastasis, and drug resistance in human malignancies, including PDAC. High expression of EGFR has been detected in the majority of cases of pancreatic cancer(Barton et al, 1991). Interestingly, the silencing of SNHG10 inhibited the activation of EGFR signaling through a significant decrease in the phosphorylation of EGFR in both SW1990 and PANC-1 cells **(Fig.5a, b)**. *KRAS* exon 12 (G12V) mutations are one of the initiating events in the tumorigenesis of PDAC and are observed in around 90% of PDAC cases(Collisson et al, 2019; Luo, 2021). The *KRAS* mutant PDAC cells have constitutive activation of its downstream targets, including AKT and ERK1/2. Furthermore, SNHG10 depletion significantly suppressed the activated AKT and ERK1/2 signaling cascade in *KRAS* mutant PDAC, whereas total AKT and ERK1/2 protein levels remained unaffected **(Fig. 5a, b)**. Several studies have reported that c-MET, hepatocyte growth factor receptor is one of the well-known oncogenic receptor tyrosine kinases which is robustly expressed and regulate tumorigenesis, cancer stemness, metastasis, and drug resistance in variety of human malignancies including pancreatic cancers through multiple signaling like mTOR, MAPK, PI3K/AKT, STAT3(De Santis et al, 2024; Qin et al, 2022; Xu et al, 2020). Interestingly, the downregulation of SNHG10 resulted in a significant reduction in the phosphorylation of c-MET and mTOR, suggesting c-MET-mediated inhibition of mTOR signalling in the SW1990 and PANC-1 cells **(Fig. 5a, b)**.

**Fig. 5.**
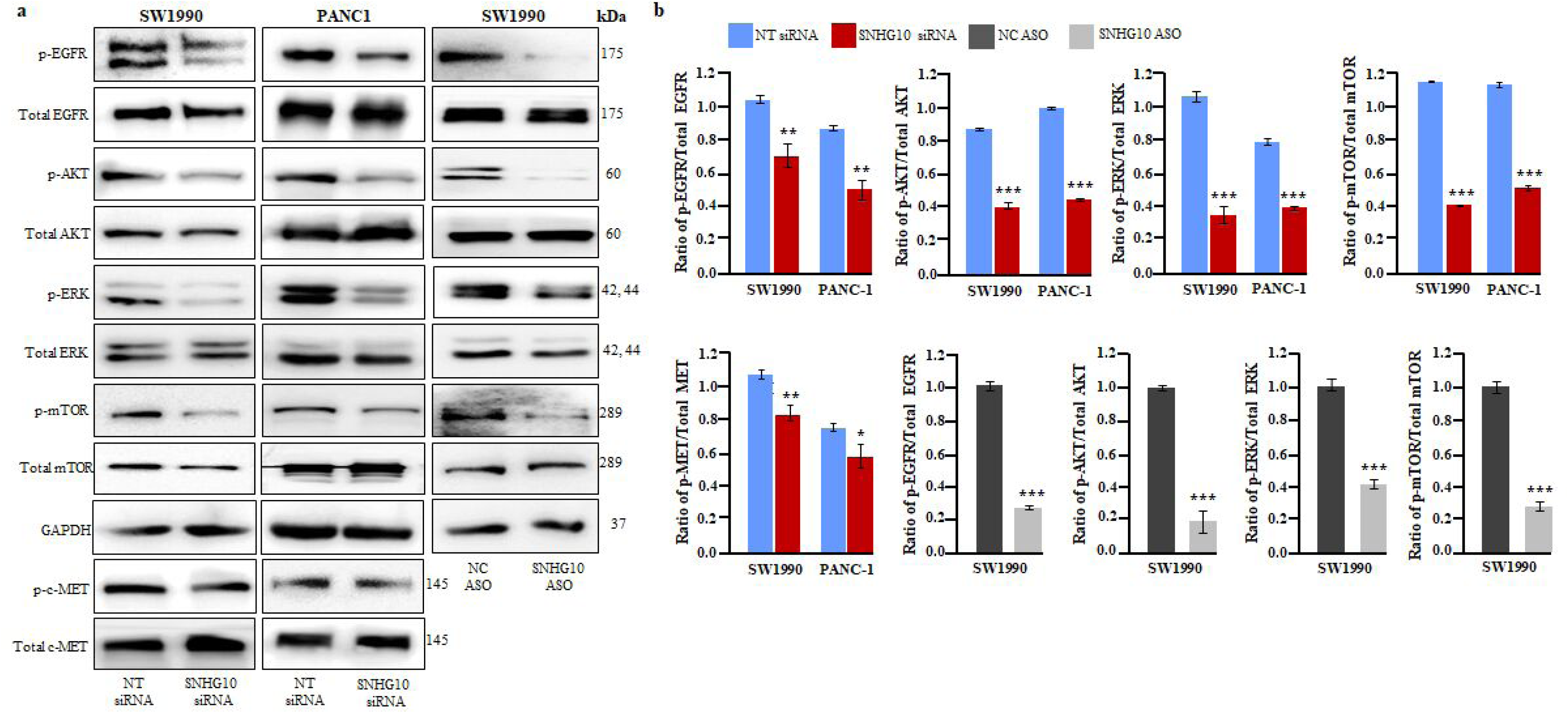
Downregulation of SNHG10 suppressed EGFR/AKT/ERK1/2 signalling cascade in PANC-1 and SW1990 cells. **(a)** The cell lysate of SNHG10-silenced PDAC cells was used for western blot analysis to detect the expression of the phosphorylation of EGFR, AKT, ERK1/2, mTOR, and c-MET. **(b)** Densitometry analysis of western blotting data by ImageJ software. Student t-test was performed for statistical significance as shown by means ± SDs; n=3, *p < 0.05, **p < 0.01, *** p < 0.001.

### Downregulation of SNHG10 enhanced the gemcitabine sensitivity in the in vitro model of gemcitabine-resistant PDAC cells

To investigate the potential role of SNHG10 in gemcitabine resistance in pancreatic cancer cells, we first generated the gemcitabine-resistant PANC-1 and CFPAC-1 cells. The PANC-1 and CFPAC-1 cells were cultured in the presence of increasing concentrations of gemcitabine. We observed a change in the morphology in the PANC-1/GemR, and CFPAC-1/Gem R cells as compared to the parental PDAC cells **(Fig. 6a; supplementary Fig. S5a).** Parental PDAC cell lines show highly colonized morphology with tight junctions and a cobblestone appearance, while gemcitabine-resistant cells showed highly invasive and migratory spindle-shaped phenotype with reduced adhesion characteristics like transformed fibroblasts **(Fig. 6a; supplementary Fig. S5a)** (Chen et al, 2022a; Samulitis et al, 2015). The PANC-1/GemR and CFPAC-1/GemR cells had significantly higher viability and lower sensitivity to Gem than parental PDAC cells. The IC50 values for PANC-1/GemR and CFPAC-1/GemR were 19.8-fold and 4-fold higher than parental, respectively **(Fig. 6b; supplementary Fig. S5b)**. Next, we also checked the expression of genes responsible for Gem resistance, including RRM1, RRM2, hENT, and hDCK. Our data displayed significantly increased expression of RRM1 and RRM2 and repressed expression of hENT in PANC-1/GemR and CFPAC-1/GemR than in parental PDAC cells **(Fig. 6c; supplementary Fig. S5c)**. Interestingly, SNHG10 expression was significantly induced in the PANC-1/GemR and CFPAC-1/GemR cells **(Fig. 6d; supplementary Fig. S5d)**. We have confirmed the significant depletion of SNHG10 expression upon treatment of SNHG10 ASO in PANC-1/GemR compared to NC ASO **(Fig. 6e; supplementary Fig. S5e)**. Furthermore, the depletion of SNHG10 in PANC-1/GemR and CFPAC-1/GemR cells restored the gemcitabine sensitivity of PANC-1/Gem and CFPAC-1/GemR cells. The increased Gem sensitivity inhibited the cell viability and clonogenic growth of the PANC-1/Gem and CFPAC-1/GemR cells **(Fig. 6f, g; supplementary Fig. S5f, g)**.

**Fig. 6.**
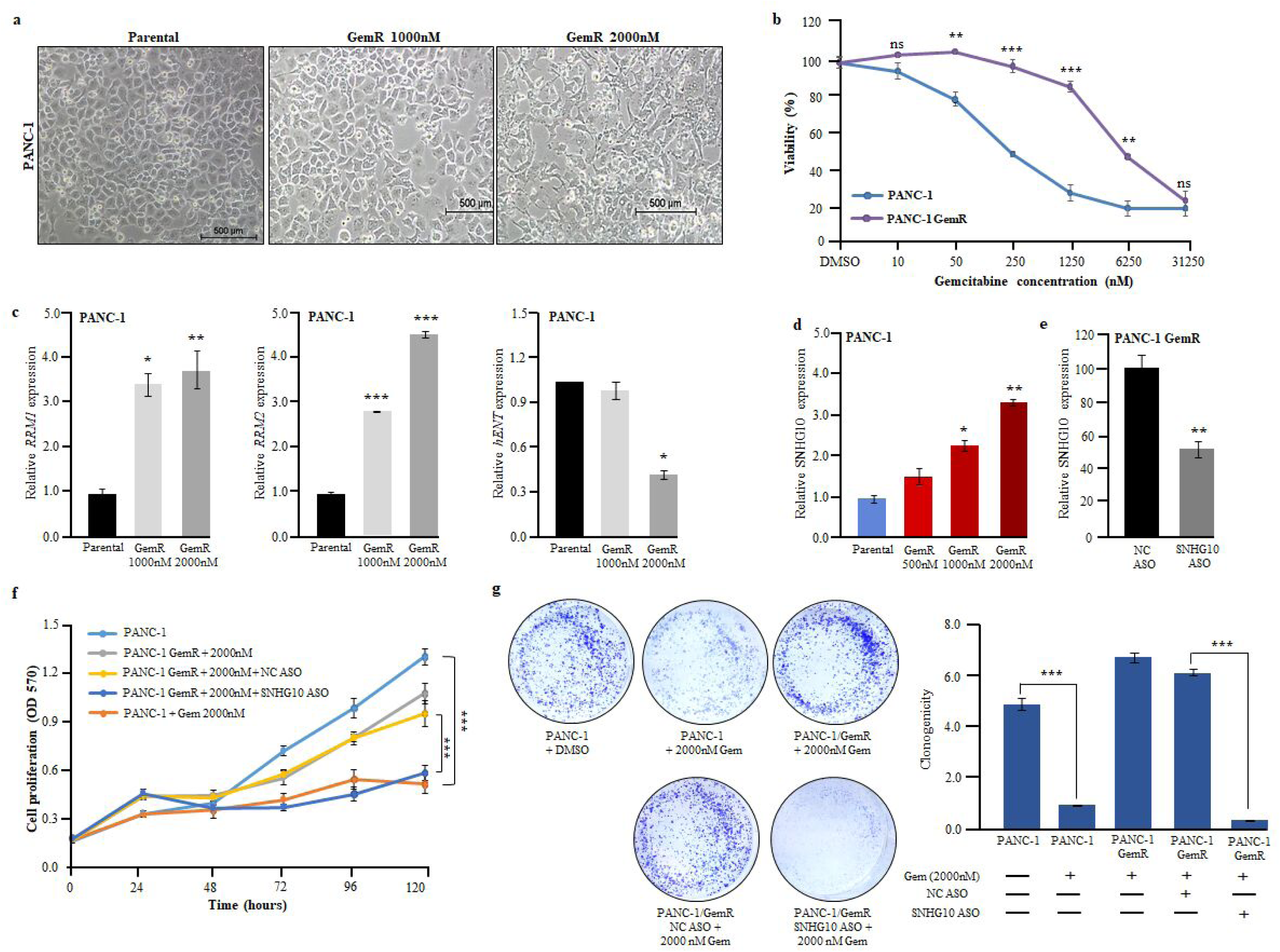
Downregulation of SNHG10 enhanced the gemcitabine resistance in the in vitro model of gemcitabine-resistant PDAC cells. **(a)** Representative images showed migratory and invasive spindle morphology in gemcitabine-resistant PANC-1 cells. (**b)** MTT assay confirmed differential sensitivity of gemcitabine. (**c)** Expression analysis of gemcitabine resistance-related genes (*RRM1*, *RRM2*, *and hENT*). (**d)** qPCR data displayed induction of SNHG10 in PANC-1 gemcitabine-resistant cells. (**e)** Knockdown of SNHG10 in gemcitabine-resistant cell line confirmed by qPCR. (**f** and **g)** The proliferation and clonogenic assays demonstrated gemcitabine sensitivity upon SNHG10 depletion.

### Efficacy of SNHG10 ASO treatment on tumor growth in PDAC xenograft model

To investigate the effectiveness of SNHG10 downregulation using SNHG10 in vivo grade ASO on tumor growth, we established a PDAC xenograft model by subcutaneous injection of SW1990 cells into the abdominal region of NOD-SCID mice. The SNHG10 ASO treatment group of SW1990 xenograft showed a significant reduction in tumor volumes and growth compared to the NC ASO group **(Fig. 7a, b).** At the end of the experiment, mice were euthanized, and tumors were excised. Further, the RNA was isolated from the xenograft tumors and subjected to qPCR analysis. The qPCR data displayed significant depletion in SNHG10 transcript levels in the SNHG10 ASO-treated xenografts compared to NC ASO-treated xenografts **(Fig. 7c)**. The average tumor weight in the SNHG10 ASO-treated xenograft group was significantly reduced than the NC ASO-treated xenograft group of mice **(Fig. 7d).** Noticeably, the SNHG10 ASO-treated xenografts revealed approximately 50% decreased tumor growth by 28 days. We performed haematoxylin and eosin (H&E) staining of SNHG10 ASO and NC ASO-treated xenograft tumor sections. The cells of SNHG10 ASO-treated xenograft tumors displayed mildly pleomorphic vesicular nuclei, whereas moderate vesicular nuclei were noted in NC ASO-treated xenograft tumors. The SNHG10 ASO-treated tumor sections showed apoptosis at the intervening stroma compared to poorly differentiated tumors in the NC ASO-treated group **(Supplementary Fig. S6a)**. The immunohistochemical analysis displayed positive staining for CA19.9, a molecular marker for pancreatic cancer **(Supplementary Fig. S6b)**. The H&E staining of the liver section of the xenograft did not show significant histological change, suggesting that ASOs were non-toxic **(Supplementary Fig. S6c)**. The qPCR results displayed significantly increased expression of *E-cadherin* and reduced expression of *N-cadherin, vimentin, fibronectin, survivin, and HIF-1α* **(Supplementary Fig. S6d).** The immunohistochemical analysis demonstrated that SNHG10 ASO-treated xenograft tumor sections showed significantly reduced expression of cyclin D1 (cell cycle), Ki-67 (proliferation marker), vimentin (mesenchymal marker), and restoration of E-cadherin expression (epithelial marker) compared with NC ASO-treated tumor sections **(Fig. 7e, f)**.

**Fig. 7.**
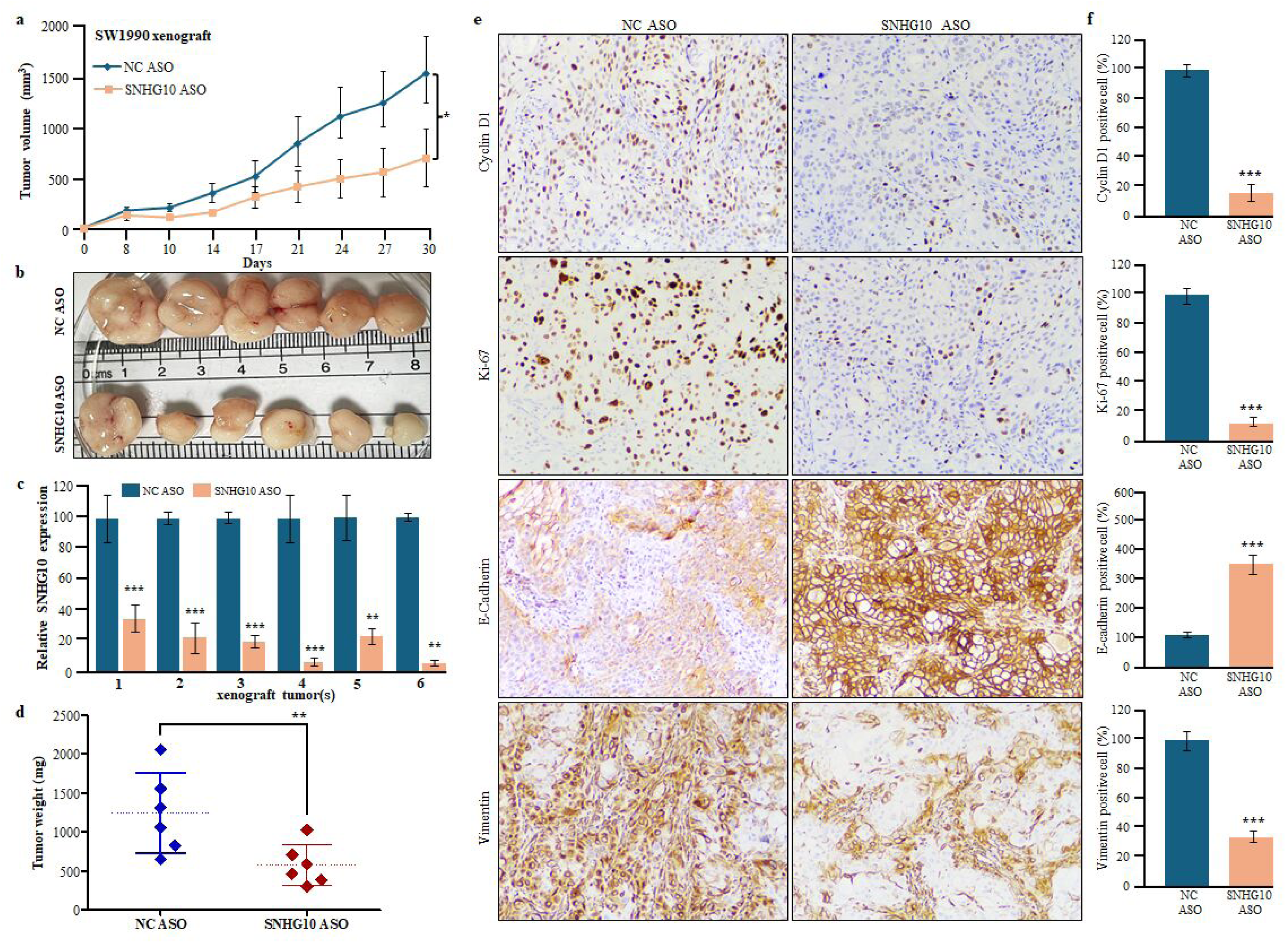
Efficacy of SNHG10 ASO treatment on tumor growth in PDAC xenograft model. **(a)** The average tumor volume between the SNHG10 ASO-treated and NC ASO-treated groups. **(b)** The photograph displayed the reduced tumor size in the SNHG10 ASO-treated group versus the NC ASO-treated group. (**c)** The qRT-PCR data revealed reduced SNHG10 expression in SNHG10 ASO-treated tumors. **(d)** Tumor weight in SNHG10 ASO-treated versus NC ASO-treated groups. **(e** and **f)** Immunohistochemical staining and analysis demonstrated cyclin D1, Ki-67, E-cadherin, and vimentin staining.

### Effect of SNHG10 ASO treatment on cell cycle, EMT proteins, as well as AKT/ERK signaling cascade in PDAC xenograft

To further understand the molecular mechanism(s) associated with the arrested tumor growth mediated through SNHG10 depletion, the expression of cell cycle, survival, and EMT regulators has been examined in xenograft tumors. The results of the western blotting and densitometric analysis revealed that SNHG10 downregulation significantly reduced the expression of vimentin, cyclin D1, CDK6, and survivin and upregulated the expression of p21, E-cadherin in SW1990 xenograft tumor specimens **(Fig. 8a, b)**. Importantly, the silencing of SNHG10 significantly suppressed the phosphorylation of AKT and ERK in SNHG10 ASO-treated tumors **(Fig. 8a, b)**.

**Fig. 8.**
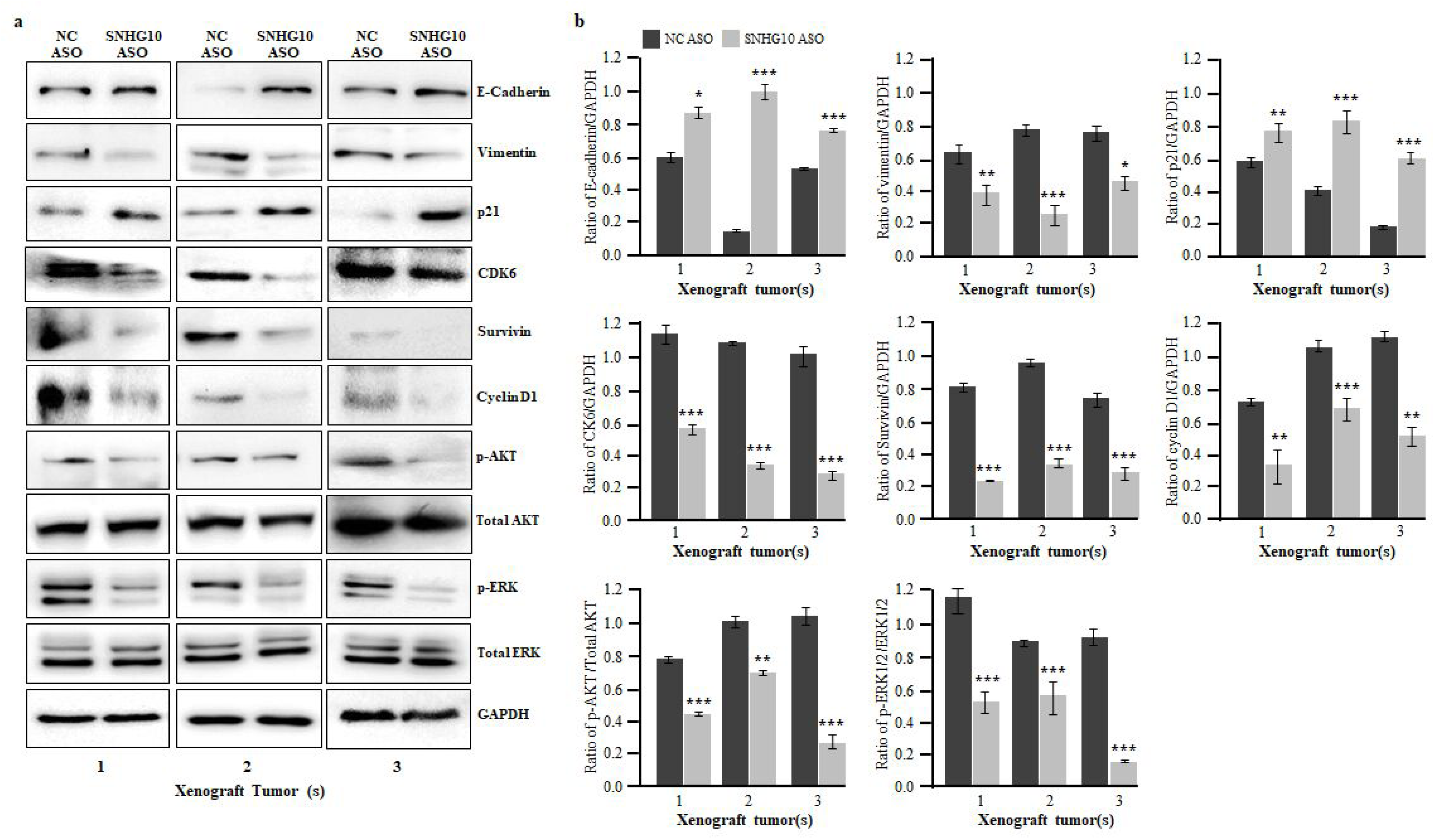
Effect of SNHG10 ASO treatment on cell cycle, EMT proteins, and AKT/ERK signaling cascade in PDAC xenograft. **(a)** The Western blot analysis on xenograft tumor samples treated with SNHG10 and NC ASO displayed the protein expression of the cell cycle and EMT regulators. The SNHG10 ASO tumors displayed inhibited AKT/ERK pathway in PDAC. (**b)** Densitometry analysis of western blotting data by ImageJ software. Student t-test was performed for statistical significance as shown by means ± SDs; n=3, *p < 0.05, **p < 0.01, *** p < 0.001.

## Discussion

Pancreatic ductal adenocarcinoma (PDAC) is a major cause of cancer-associated mortality and is often diagnosed at an advanced stage(Kirtonia et al, 2025). To date, the early diagnosis at a curable stage is mostly difficult since patients rarely show symptoms, and the non-availability of specific and sensitive biomarkers for early diagnosis of PDAC. Therefore, identifying critical molecular targets and dissecting their molecular mechanisms linked with tumorigenesis and drug resistance is essential for effective treatment of pancreatic carcinoma. Several studies have demonstrated that lncRNA plays a variety of functions in regulating transcription, translation, epigenetic, post-transcriptional, and post-translational modifications(Pandya et al, 2020; Yadav et al, 2021). Deregulation of lncRNA is associated with cancer, cardiovascular, neurological, and metabolic disease conditions(Statello et al, 2021). The lncRNA(s) have been reported to modulate cell proliferation, viability, migration, stemness, differentiation, invasion, and tumor microenvironment during cancer initiation, development, and metastasis(Kanojia et al, 2022; Shabna et al, 2023). Interestingly, the high specificity and ease of detection of lncRNA(s) in the tissues, blood, urine, and saliva have increased the interest in exploring the potential of lncRNA in human malignancies(Statello et al, 2021). SNHG10 is an important molecular target due to its robust expression in cancers(Zhu et al, 2025). To our knowledge, our study comprehensively uncovers the potential role and molecular mechanism(s) of SNHG10 in pancreatic carcinoma.

The current study demonstrated the high expression of SNHG10 in TCGA RNA sequencing data of 171 normal pancreatic samples and 179 human pancreatic ductal adenocarcinoma tissue samples, corresponding to the elevated SNHG10 expression in a panel of PDAC cell lines. The overexpression of SNHG10 was observed to be positively associated with pathological stages of patients with PDAC.

Next, we depleted the expression of SNHG10 using siRNA and antisense oligonucleotides to understand the functional relevance of SNHG10 in PDAC cell lines through phenotypic assays. Downregulation of SNHG10 transcript levels decreased cell viability and clonogenic growth of PDAC cells in liquid culture. The treatment of SNHG10 ASOs significantly suppressed tumor growth, volume, and tumor weight in the murine PDAC xenograft model, indicating that SNHG10 expression is important in the progression of PDAC. Emerging studies have confirmed that the EMT is an important cellular phenomenon, which is critical for cancer cell metastasis to distant organs, chemotherapeutic resistance, and recurrence(Lambert et al, 2017). Our data displayed that SNHG10 silencing significantly inhibited the migration through reversal EMT phenotype by restoring the E-cadherin expression (epithelial marker) with simultaneous decreased expression of N-cadherin and Vimentin (mesenchymal markers).

This study may lead to new therapeutic opportunities by regulating the cell cycle and apoptosis through SNHG10 in PDAC. The cyclinD1/CDK4/CDK6 complex performs several functions as an oncogene, by increasing multiple processes during malignant transformation, including abnormal cell cycle, cellular growth, angiogenesis, and resistance to apoptosis. SNHG10 knockdown arrested the cell cycle progression of PANC1 and SW1990 cells in different phases by inhibiting the expression of cell cycle regulator cyclin D1 and its associated kinases CDK4 and CDK6, as well as enhancing the expression of cell cycle inhibitor p21. Also, the SNHG10 depletion caused significant apoptosis(Tashiro et al, 2007). Survivin is a member of the inhibitor of apoptosis protein family that inhibits the activation of caspase protein and inhibits cell death. Survivin is robustly expressed and associated with poor clinical outcomes in pancreatic cancer. Remarkably, silencing of SNHG10 reduced the survivin in PDAC cell lines. Aurora kinases A and B, the two serine/threonine-protein kinases associated with spindle formation, centrosome, and spindle orientation during mitosis, are known for their involvement in apoptosis, metastasis, and tumorigenesis. In this context, the downregulation of SNHG10 levels decreased the Aurora A and B expression in PDAC cells.

Now, lncRNAs are known to modify gene expression at multiple levels, including transcriptional, epigenetic, and translational mechanisms. The lncRNA can interact with RNA, DNA, and proteins to modulate chromatin structure, recruit transcription factors, and regulate mRNA stability or translation. Also, lncRNA acts as a molecular sponge for miRNA, preventing them from inhibiting the expression of their target genes. To check the miRNA target of SNGH10, we performed the bioinformatic analysis with the help of Starbase, lncRNASNP2, LncBase, and miRcode, which predicted miR-150-5p with 7-mer similarity in the seed sequence of SNGH10. The downregulation of miR-150-5p in the PDAC patients and cell lines. Further, the in-silico analysis predicted VEGF-A as one of the most important targets of miR-150-5p. Importantly, the Ago2 RNA immunoprecipitation data confirm the physical interaction among SNHG10, miR-150-5p, and VEGF-A at the RISC complex. Interestingly, the depletion of SNHG10 restored miR-150-5p expression, leading to repression of VEGF-A expression in PDAC cells. These data indicated that miR-150-5p behaves as a tumor suppressor, which is consistent with other studies where miR-150-5p was downregulated in colorectal carcinoma, choriocarcinoma, and osteosarcoma through the VEGFA/PI3K/AKT/mTOR axis(Chen et al, 2018; Liao et al, 2024; Qin et al, 2020).

To uncover the molecular mechanisms and downstream signaling cascades modulated by the SNHG10/miR-150-5p/VEGF-A axis in PDAC, we have checked the effect of SNHG10 silencing on EGFR/AKT/ERK/mTOR signaling cascade. Our data indicated that downregulation of SNHG10 suppressed the activation of the EGFR/AKT/ERK//mTOR signaling cascades in PDAC cells. Recently, studies supported that the HGF/c-Met signaling pathway can activate the mTOR/NGF axis in pancreatic carcinoma(Qin et al, 2022). The downregulation of mTOR may be partially due to the inhibition of c-MET signaling upon SNHG10 silencing.

Chemotherapeutic drug resistance is the major challenge in pancreatic carcinoma treatment in clinics. Furthermore, unravelling the effect of SNHG10 on the mechanism of gemcitabine resistance is important for optimizing current therapeutic strategies. Interestingly, the SNHG10 ASO treatment restored the gemcitabine sensitivity in PDAC gemcitabine-resistant cells, which resulted in significant decreased cell growth and clonogenic ability.

Altogether, the findings of this study confirmed the oncogenic potential of SNHG10 and its role in the gemcitabine resistance in PDAC cells through modulation of molecular targets directly or indirectly at the transcription and translation level.

In summary, this study highlights that SNHG10 holds great potential for diagnostic and therapeutic targeting of the SNHG10-mediated miR-150-5p/VEGF-A/EGFR/AKT/ERK/mTOR axis **(Fig. 9)** for the management of tumorigenesis and gemcitabine resistance of PDAC.

## Materials and Methods

### Cell culture

AsPC-1, BxPC-3, Capan-2, PANC-1, Panc10.05, SW1990, MIA PaCa-2, and CFPAC-1 were purchased from the American Type Culture Collection (ATCC, USA). AsPC1, BxPC-3, and Panc10.05 were grown in RPMI-1640 (Gibco, USA); MIA PaCa-2 and PANC-1 were maintained in DMEM (Gibco, USA)(Kirtonia et al, 2022). Leibovitz’s L-15, McCoy’s 5A, and Iscove’s modified medium (Gibco, USA) were used to culture SW1990, Capan-2, and CFPAC-1, respectively(Kirtonia et al, 2022). The media were supplemented with 10% FBS (Gibco, USA) and 1% penicillin-streptomycin (Invitrogen, Carlsbad, CA), and cells were grown in a 37^0^C incubator with 5% CO_2_ and controlled humidified conditions. Human normal pancreatic epithelial (hTERT-HPNE) cells were grown using a specific media having 25% M3 medium (INCELL, Texas, USA), 75% DMEM (Gibco, USA), puromycin (Sigma-Aldrich, Merck, Germany), and 10ng/ ml EGF (Thermo-Fisher Scientific, USA).

### RNA extraction, complementary DNA (cDNA) synthesis, and quantitative real-time PCR (qRT-PCR) analysis

Total RNA was isolated from AsPC-1, BxPC-3, Capan-2, PANC-1, Panc10.05, SW1990, MIA PaCa-2, and CFPAC-1 and hTERT-HPNE cells using the RNeasy mini kit (Qiagen, GmbH, Germany) and TRI Reagent (Ambion Inc., Austin, TX) as described previously(Deswal et al, 2024; Kanojia et al, 2013). Total RNA was treated with DNase I to avoid genomic DNA contamination before cDNA synthesis. 2μg of total RNA was used for cDNA using iScript reverse transcription Supermix (Bio-Rad, USA) specifically for the expression of lncRNA. The miRNA-specific cDNAs were prepared using specific stem-loop primers and high-capacity cDNA reverse transcription kit (Thermo-Fisher Scientific, USA). The qRT-PCR was performed using Powerup SYBR Green PCR Master Mix kit (Thermo-Fisher Scientific, USA) using the ABI StepOnePlus real-time PCR System (Thermo-Fisher Scientific, USA) for both lncRNA and miRNA. The *U6* and *GAPDH* transcripts were used as endogenous controls. The delta threshold value (ΔCt) was calculated from the given threshold (Ct) value using the formula ΔCt = (Ct *SNHG10* – Ct *U6* or *GAPDH*) for each sample. The sequences of the primers are mentioned in **Supplementary Table 1**.

### Transfection of small-interference RNA (siRNA) and antisense oligonucleotide (ASO)

SNHG10-specific siRNA (SNHG10 siRNA), and non-target (NT siRNA) were designed and purchased from Dharmacon, Horizon Discovery, USA, through Biogene India(Kanojia et al, 2017). SNHG10 and negative control (NC) ASOs were designed using the Qiagen online tool for both *in vitro* and *in vivo* experiments. Transfection was performed in PDAC cell lines with Lipofectamine 3000 reagent (Invitrogen, Thermo-Fisher Scientific, USA) according to the protocol as provided by the manufacturer. The sequences of SNHG10 siRNAs and ASOs are provided in **Supplementary Table 2**.

### Cell viability assay

Methyl thiazolyl-diphenyl-tetrazolium bromide (MTT; Sigma-Aldrich, Merck, USA) assay(Shahid et al, 2024) was performed to assess the effect of SNHG10 knockdown on the cell viability and growth of PDAC cells as described earlier(Kirtonia et al, 2025). Briefly, 4000 cells per well were seeded in 96-well plates (Cole-Parmer) in 100 μl of medium and incubated in a CO_2_ incubator at 37°C. After 24 hours, transfection was performed using SNHG10 siRNA/ASO and NT siRNA/NC ASO. The culture medium was replaced with a 100 μl serum-free medium containing 0.5mg/ml MTT and incubated for 3 hours in a CO2 incubator at 37°C at indicated time point (0, 24, 48, 72, 96 hours) of transfection. Formazan crystals were dissolved in 100 uL of DMSO. Absorbance was measured at 570 nm using Multiskan GO spectrophotometer (Thermo-Fisher Scientific, USA).

### Colony formation assay

To study the effect of SNHG10 knockdown on the clonogenic ability, 500 cells were seeded into 12-well plates in triplicate in a complete medium, and transfection was performed using siRNA/ASO twice a week. After 2 weeks, colonies were visible, fixed with cold methanol, stained with 0.1% crystal violet, and photographs were captured. The stained colonies were dissolved in DMSO, and absorbance was measured at 570 nm using Multiskan GO (Thermo-Fisher Scientific, USA) spectrophotometer.

### Cell cycle assay

Control cells and SNHG10 knockdown cells were harvested after 48 hours for transfection and washed thrice with 1X PBS (cold). These cells were fixed with 75% ethanol (pre-cooled at −20°C) at 4°C overnight and washed with 1X PBS followed by staining with 500 μl of propidium iodide staining solution (50 μg/ml PI, 0.1%Triton X-100 (v/v), 20 μg/ml DNase-free RNase in PBS) for 30 minutes at room temperature in the dark(Gala et al, 2024; Garg et al, 2017). Ten thousand events per sample were acquired using a BD ARIA-III flow cytometer (BD, CA, USA), and the percentage of cells in G0/G1, S, G2/M, and Sub-G2/M phases of the cell cycle was examined with FACS Diva software.

### Apoptosis assay

To evaluate the effect of SNHG10 knockdown on apoptosis, SNHG10 siRNA/ASO and NT siRNA/NC ASO transfected cells were harvested after 48 hours of transfection and washed with 1X cold PBS at least two to three times. After washing, cells were stained with Annexin V-FITC and PI using an Apoptosis Detection Kit (Thermo-Fisher Scientific, USA) as per manufacturer protocol at room temperature for 30 minutes(Hayano et al, 2013). Then cells were analyzed using a BD ARIA-III flow cytometer (BD, CA, USA), and ten thousand events were recorded.

### Cell migration and invasion assay

1×10^5^, SNHG10 depleted cells and control cells were suspended in 200µl FBS-free medium and placed in the top chamber of a transwell insert (8-μm pore size) in triplicate. The lower chamber was filled with 400μl medium containing 10% FBS as a chemoattractant and was incubated in a humidified CO2 incubator at 37 °C. After 48 hours, cells from the top chamber were removed using a cotton swab, and cells on the lower surface of the insert were fixed using chilled methanol and stained with 0.1% crystal violet. Cells were counted using a bright-field microscope.

### Western blotting

SW1990 and PANC-1 cells transfected with SNHG10 specific siRNA/ASO, NT siRNA, and NC ASO were harvested after 48 hours of transfection and lysed using Mammalian protein extraction buffer (Thermo-Fisher Scientific, USA) followed by protein estimation as described previously(Garg et al, 2015; Sneha et al, 2020). Briefly, 30 ug of proteins were loaded and separated on 10% SDS-PAGE followed by transfer to a PVDF membrane (Millipore, Merck). The membranes were blocked with 5% fat-free milk for 30 to 40 minutes at room temperature, then incubated overnight at 4°C with specific primary antibodies (1:1000 dilution) followed by 3 times washing with PBST. Next, membranes were incubated with the HRP-conjugated secondary antibody for 2 hours at 1:5000 dilution. The protein signals were visualized with enhanced chemiluminescence (Thermo-Fisher Scientific, USA). GAPDH antibody was used for equal loading control. The details of the primary antibodies are provided in **Supplementary Table 3**.

### RNA Immunoprecipitation Assay (RIP)

To perform the RIP assay, 2×10^7^ PDAC cells were trypsinsed from two 100 mm dishes. PDAC cell pellet was washed with cold PBS and further lysed by adding 200µL of Mammalian protein extraction reagent. The supernatant of the lysed PDAC cells was incubated with anti-Ago2 antibody (CST, USA), and normal IgG (CST, USA) was added in a 1:50 dilution. Protein-A agarose Fast Flow beads (Millipore, USA were washed with 1XPBS, and the beads pellets were saved. Further, the beads were added to the cell lysate and incubated in the tubes on a rotator at 10 RPM overnight at 4^0^C. The next day, beads were washed with MPER, and coprecipitated RNAs were isolated using Trizol reagent (Ambion by Life Technologies), cDNA was synthesized using high-capacity cDNA synthesis Reverse Transcription kit (Applied Biosystem, Thermo-Fisher Scientific, USA), and qRT-PCR was performed to assess the enrichment of SNHG10, miR-150-5p, and VEGF-A.

### Generation of gemcitabine-resistant human pancreatic cancer cell lines

The gemcitabine-resistant PANC-1 and CFPAC-1 cell lines were generated by exposing PANC-1 and CFPAC-1 cells to increasing concentrations of gemcitabine. The PANC-1 and CFPAC-1 cells were initially treated with 0.1 to .5 µg/ml for two weeks. The median lethal doses of gemcitabine treated PANC-1 and CFPAC-1 cells were determined by MTT assay. Then, PANC-1 and CFPAC-1 cells were cultured in 1 µg/ml gemcitabine for 72 hours. The medium containing dead cells was discarded. The fresh drug-free medium was added, and cells were passaged after attaining the confluency. Again, the cells were maintained at 1 µg/ml for two months. Next, the cells were cultured in 2 to 5µg/ml of gemcitabine for another 2 months. After 8 months, we were successful in generating gemcitabine-resistant cell lines. Resistant cell lines were labelled as PANC-1 GemR and CFPAC-1 GemR.

### PDAC xenograft model

All the mice xenograft experiments were approved by the Institutional Animal Ethical Committee of the Regional Centre for Biotechnology (RCB), Faridabad (Approval no. RCB/IAEC/2022/145). The mice experiments were performed using the best veterinary practices as reported earlier(Arora et al, 2021). A total of 12 NOD-SCID female mice at ages 8-9 weeks were randomly divided into two groups of six mice each(Garg et al, 2009). SNHG10 specific and negative control (NC) ASO were designed for in vivo delivery in xenograft murine models. 2×10^6^ SW1990 cells were resuspended in RPMI medium and Matrigel in 1:1 dilution and injected subcutaneously in the abdomen region of the mice. We allowed the SW1990 cells to form measurable tumors within the volume ranges of 50 to 100 mm^3^ for 10 days and randomly divided into two treatment groups: (i) SNHG10 ASO and (ii) NC ASO. After this, we started the administration of SNHG10 ASO and NC ASO at a concentration of 100nM in 100μL of RPMI medium into the peritoneum of SNHG10 and NC murine groups twice a week for 4 weeks, respectively(Gutschner et al, 2013; Li et al, 2020; Littlejohn et al, 2008). The volumes of the tumors were measured with a vernier caliper in both groups before the administration of ASO. The growth of the tumors was monitored for a total of 30 days. All the mice were euthanized as per the guidelines of the Institutional Animal Care and Use Committee (IACUC), one week after the last injection. The tumors were harvested, snap-frozen in liquid nitrogen, and stored at −80^0^C for qRT-PCR and immunohistochemistry experiments(Li et al, 2022).

### Immunohistochemistry (IHC) assay

The IHC was performed on the xenograft tumor tissue sections as described previously(Kirtonia et al, 2022). Briefly, the paraffin-embedded xenograft tumor sections were deparaffinized with xylene for 10 minutes at least thrice and hydrated using graded ethanol. Antigen retrieval was achieved using we used citrate buffer (pH 6.0) for antigen retrieval under steam for 15 to 20 min followed by peroxidase blocking. The indicated primary antibodies were used in DAKO protein blocking in a 1:200 dilution. The slides were developed as per the manufacturer’s instructions using diaminobenzidine followed by haematoxylin staining, and mounted with DPX (Sigma-Aldrich, St. Louis, MO).

### Statistical analysis

GraphPad Prism (San Diego, CA) and Student t-test were applied to calculate the statistical significance of the experiments. The results were displayed as mean ± standard deviation. All experiments were performed in biological triplicate and a minimum of three times.

## Supplementary Figure Legends

**Supplementary Fig. S1.**
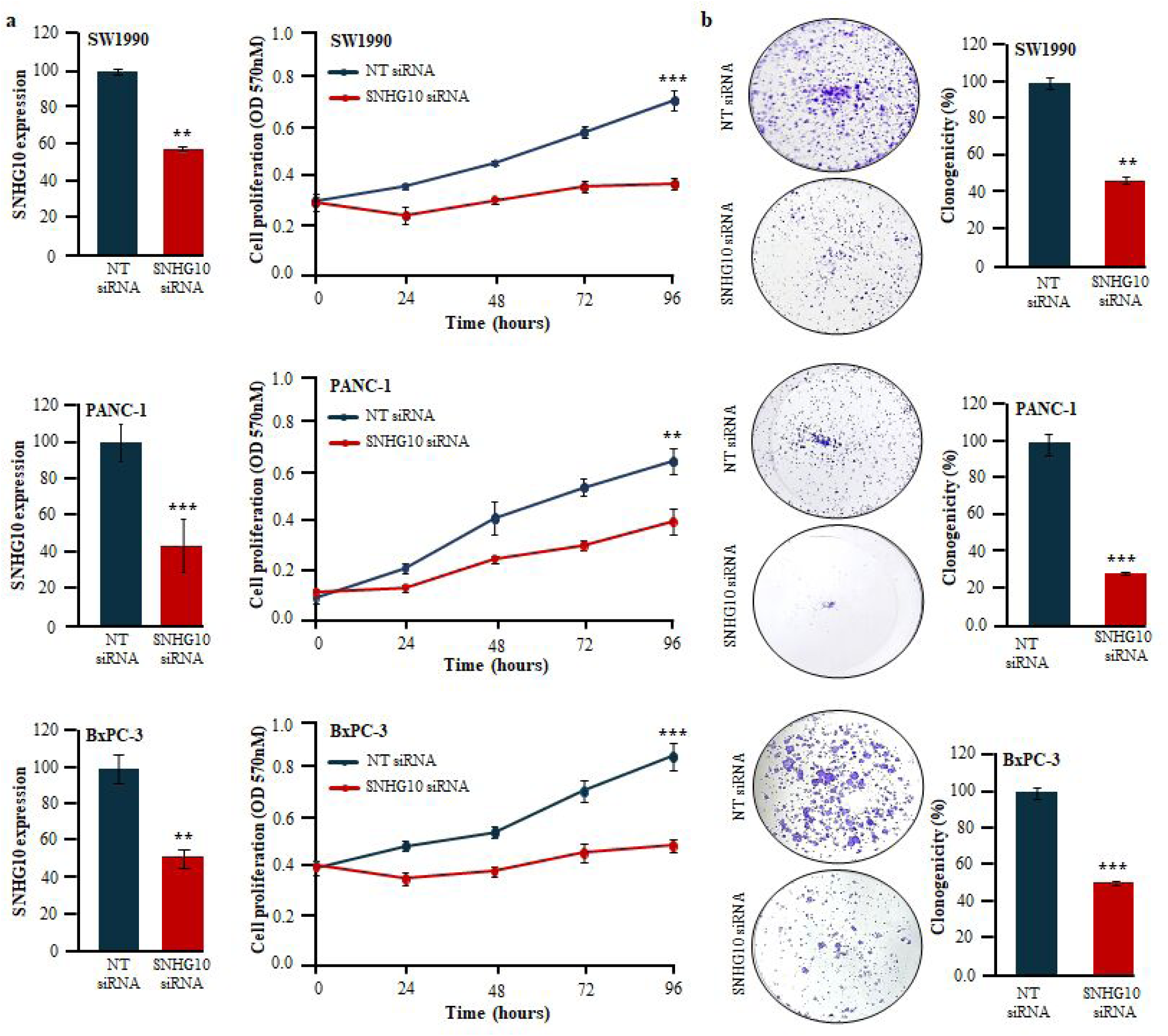
Depletion of SNHG10 significantly suppresses the viability and clonogenicity of PDAC cells. **(a)** The qRT-PCR displayed the depletion of SNHG10 in siRNA-transfected SW1990, PANC-1, and BxPC-3 cell lines. MTT assay measures the proliferation ability of SW1990, PANC-1, and BxPC-3 cells transfected with SNHG10 siRNA. **(b)** Colony formation assay confirmed clonogenicity of SNHG10 siRNA-transfected PDAC cells. All the experiments were performed three times in biological triplicate. The results are presented as the means ± SDs; n=3. *p < 0.05, **p < 0.01, ***p < 0.001; two-tailed Student t test.

**Supplementary Fig. S2.**
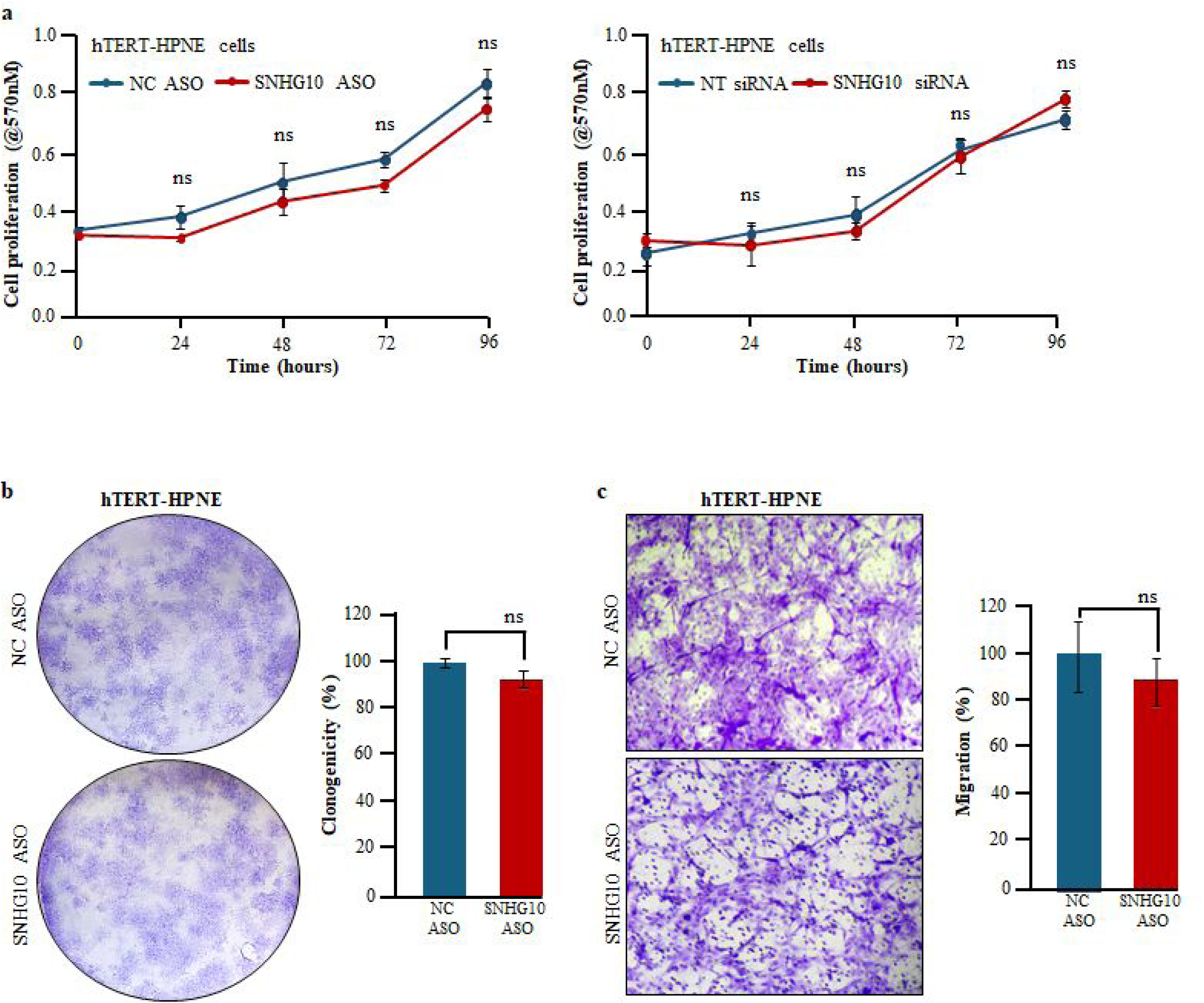
Effect of SNHG10 depletion on cell viability, clonogenicity, and migration of human normal pancreatic epithelial cells. **(a)** The cell viability was confirmed by MTT assay. **(b)** Clonogenicity measured the colony-forming capacity of hTERT-HPNE. **c** Boyden chamber assay determined the migration ability of hTERT-HPNE cells. Student t-test was performed for statistical significance. ns; not significant.

**Supplementary Fig. S3.**
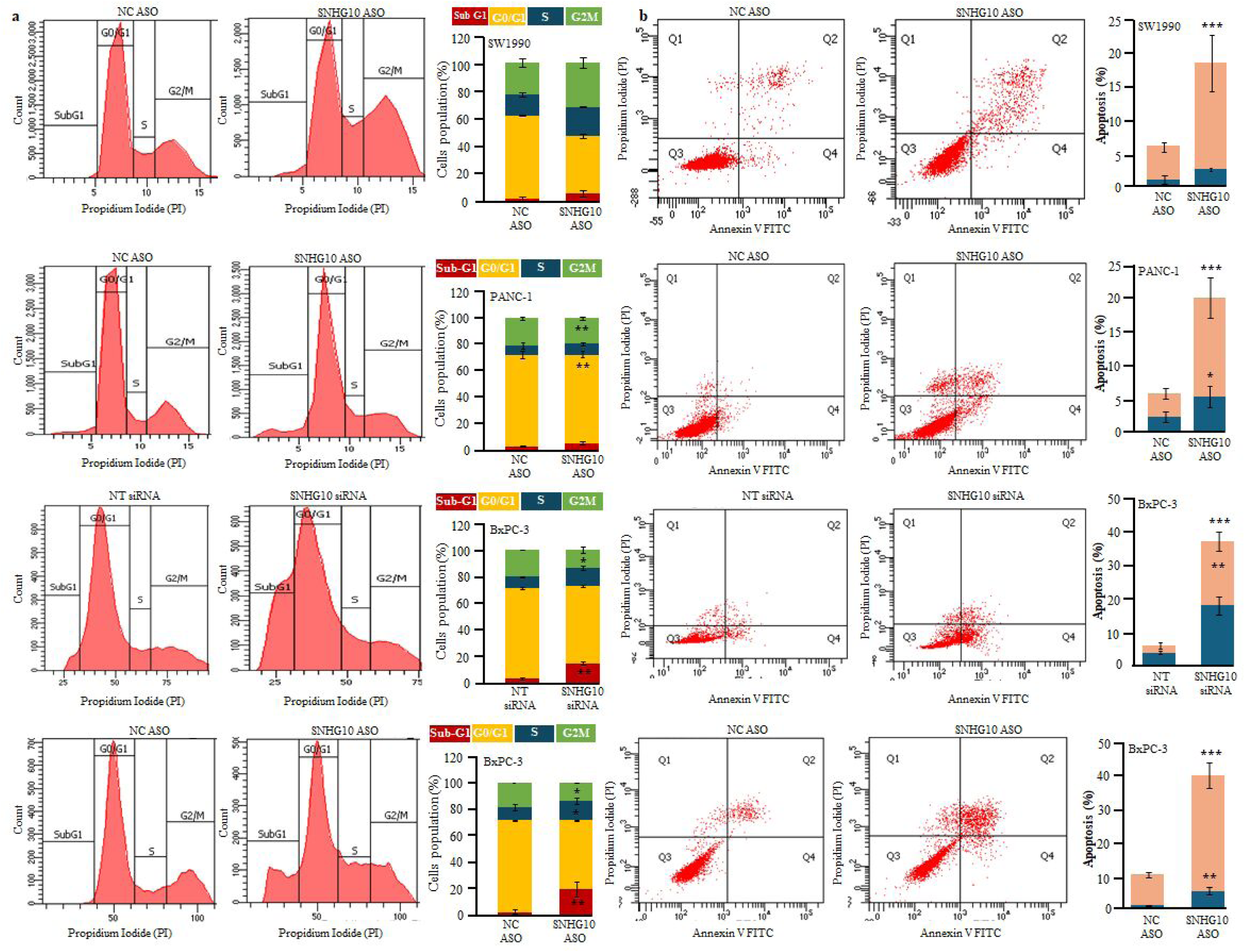
SNHG10 depletion induced cell cycle arrest and apoptosis. **(a)** Cell cycle distribution measured by propidium iodide staining in SNHG10-depleted SW1990, PANC-1, and BxPC-3 cells. Quantification and histogram represented the cell population in different phases of the cell cycle. (**b)** Cell apoptosis was determined using Annexin V and propidium iodide staining in SNHG10-depleted SW1990, PANC-1, and BxPC-3 cells.

**Supplementary Fig. S4.**
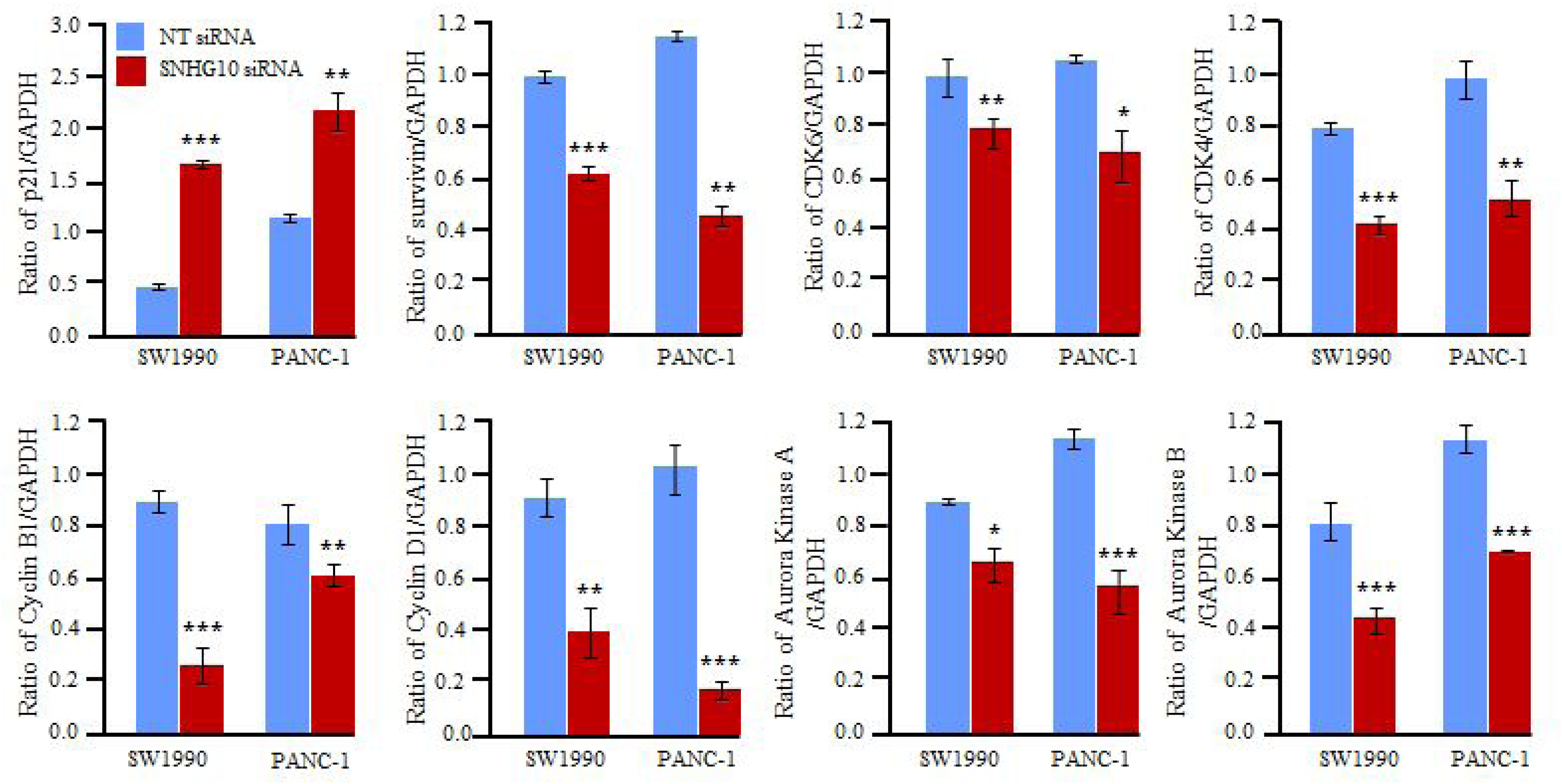
Densitometric analysis of cell cycle and apoptosis regulators in PDAC cells. Densitometry of Western blots of SNHG10-silenced PDAC cell lines displayed a significant difference in the expression of p21, survivin, CDK4, CDK6, cyclin B1, cyclin D1, arora kinase A, and arora kinase B proteins.

**Supplementary Fig. S5.**
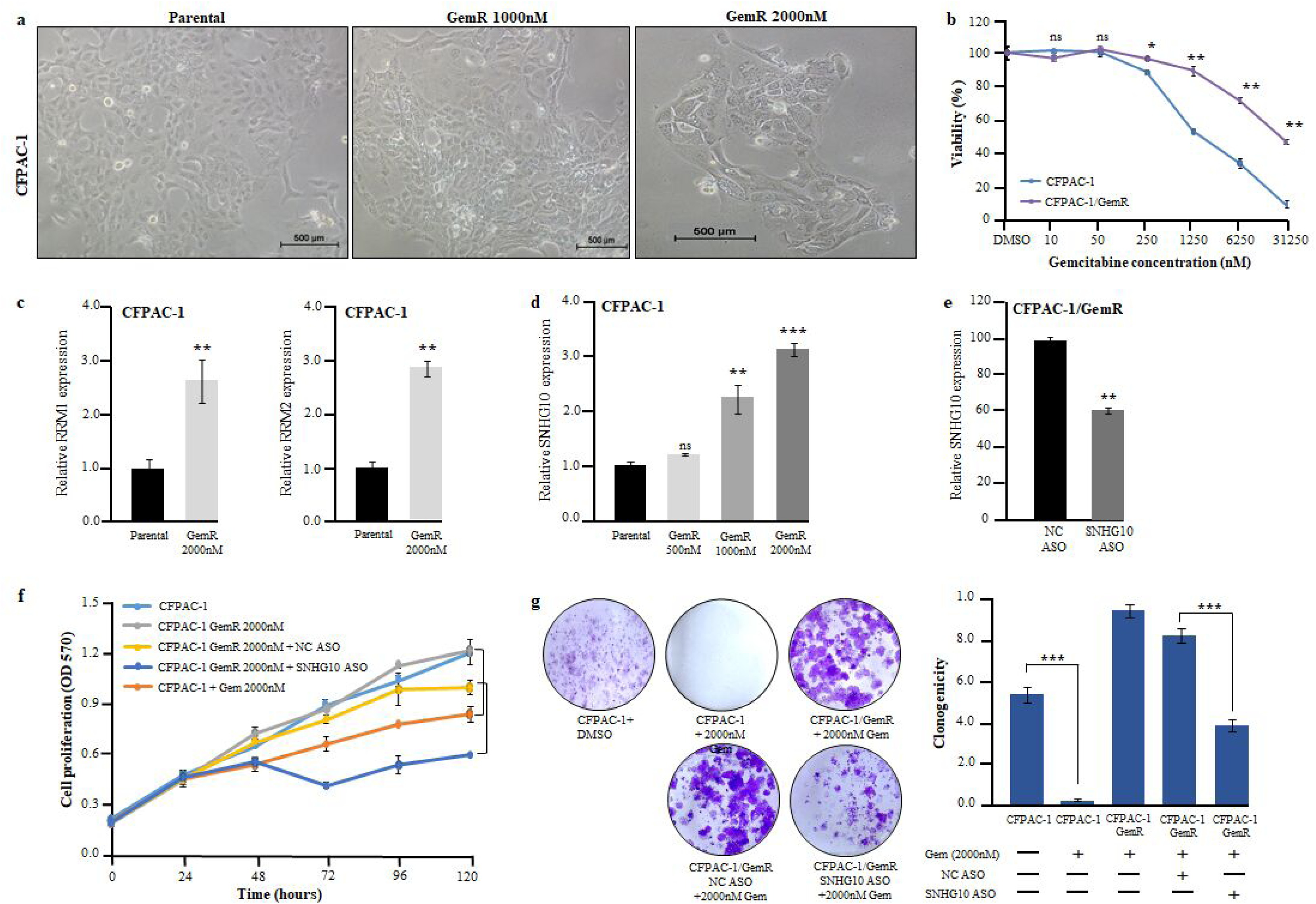
Downregulation of SNHG10 enhanced the gemcitabine resistance in the in vitro model of gemcitabine-resistant PDAC cells. **(a)** Representative images showed morphological changes in gemcitabine-resistant CFPAC-1 cells. **(b)** MTT confirmed the differential sensitivity of gemcitabine. **(c)** Expression analysis of gemcitabine resistance-related genes. **(d)** PCR data displayed the induction of SNHG10 in gemcitabine-resistant cells. **(e)** qPCR displayed knockdown of SNHG10. **(f, g)** Proliferation and clonogenic assays demonstrated gemcitabine sensitivity.

**Supplementary Fig. S6.**
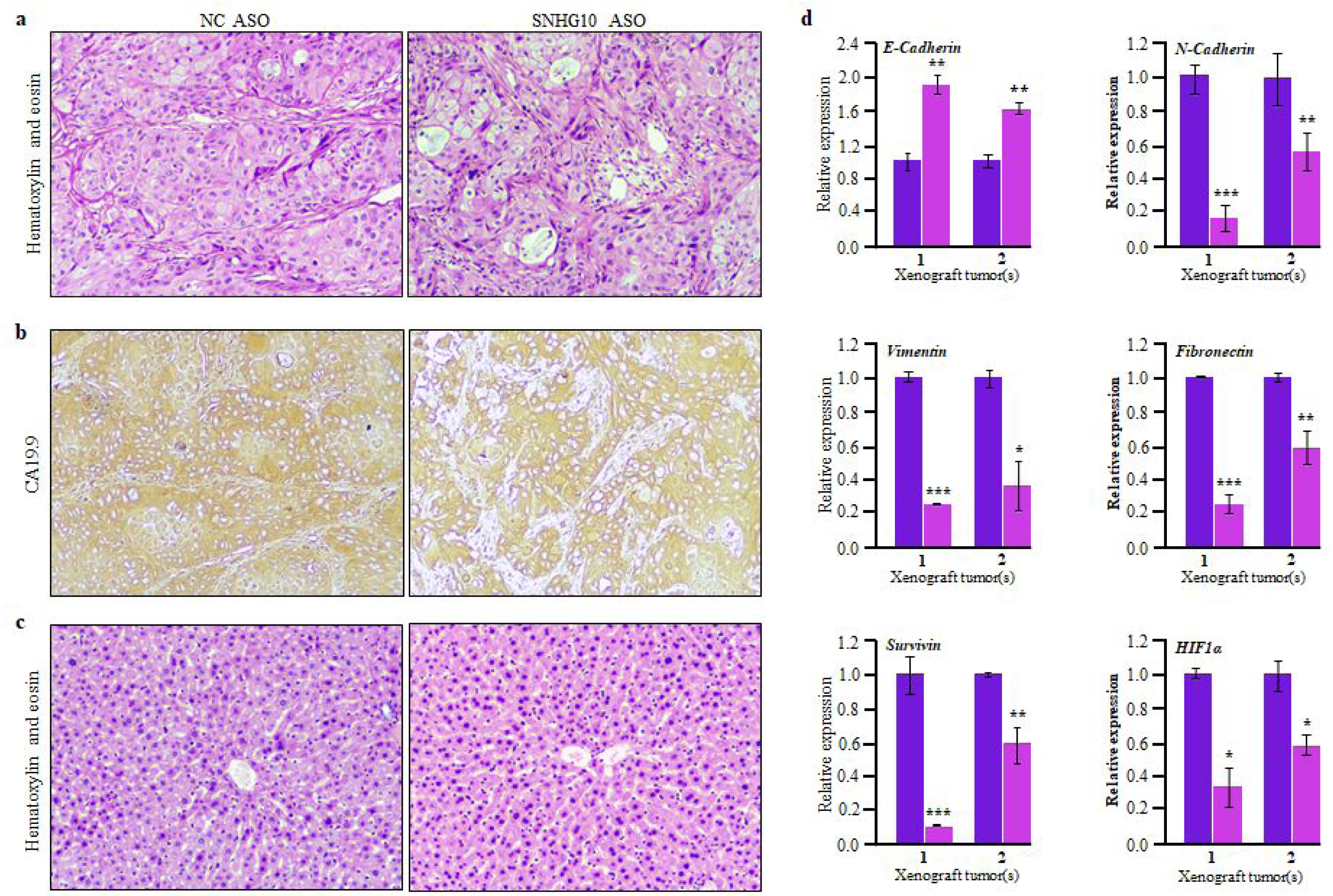
Effect of SNHG10 ASO treatment in PDAC xenograft model. **(a-c)** H&E, and IHC staining of SNHG10 ASO-treated and NC ASO xenograft tumor and liver sections. **(b)** CA19.9 staining confirms the origin of PDAC. **(c)** The qRT-PCR data revealed *E-cadherin*, *N-cadherin*, *vimentin*, *fibronectin*, *survivin*, and *HIF-1α* expression in ASO-treated tumors.

## DATA AVAILABILITY

The TCGA RNA sequencing datasets used for analysis are available at the TCGA portal. Original uncropped blots are provided in the supplementary information.

## ACKNOWLEDGMENTS

We acknowledge the Department of Biotechnology (DBT), Government of India, for the Ramalingaswami Fellowship (BT/RLF/Re-entry/24/2014) awarded to Prof. Manoj Garg. This work was supported by the Indian Council of Medical Research (ICMR), the Government of India, under Extramural Ad-hoc grants (No. 5/13/6/2022/NCD-III; 2021-12187). We also acknowledge the Department of Science and Technology (DST) and Amity University Uttar Pradesh for supporting the FIST-Flow Cytometry Facility (SR/FST/LS-II/2017/115) at the Amity Institute of Molecular Medicine and Stem Cell Research. We acknowledge the support of the Council of Scientific and Industrial Research (CSIR) and the University Grants Commission (UGC) for awarding the Senior Research Fellowship to Miss Gouri Pandya (Ref. No. 932). We thank the ICMR (No. 5/13/6/2022/NCD-III; 2021-12187) for supporting Miss Aishwarya Singh as Research Scientist-1 under the guidance of Prof. Manoj Garg. Rajender K Motiani acknowledges the funding support from the DST-Science and Engineering Research Board funding (CRG/2023/004054). Deepti K Pandita and Manoj Garg acknowledge the funding support from the Anusandhan National Research Foundation (ANRF), SERB funding (CRG/2023/003482).

## AUTHOR CONTRIBUTIONS

Manoj Garg conceived the idea, designed the study, reviewed, edited, validated, supervised, and acquired funding support. Gouri Pandya, Aishwarya Singh, Rachana Kumari, Rashi Sharma performed the experiments, acquired and analyzed the data. Suman Sourav, Sharon Raju, Rajender K Motiani, Deepti Pandita, and Manoj Garg have designed and performed xenograft experiments. Gouri Pandya and Manoj Garg prepared the initial draft of the manuscript. Gautam Sethi, Amit Kumar Pandey, Rajender K Motiani, Bhudev C Das, Deepti Pandita, and Manoj Garg reviewed, edited, and finalized the manuscript and figures for submission. All the authors have read and approved the final manuscript.

## ADDITIONAL INFORMATION

Supplementary information is available under the supplementary material.

### Consent for publication

The authors have agreed and given their consent to publish this manuscript.

### Ethics approval and consent to participate

All the experiments were carried out at the research facilities available at Amity University Uttar Pradesh, Noida, as per the approval of the Institutional Biosafety Committee. Animal experiments were carried out with prior approval (Approval number RCB/IAEC/2022/145) from the Institutional Animal Ethics Committee of the Regional Centre for Biotechnology, Faridabad, Haryana, India.

---

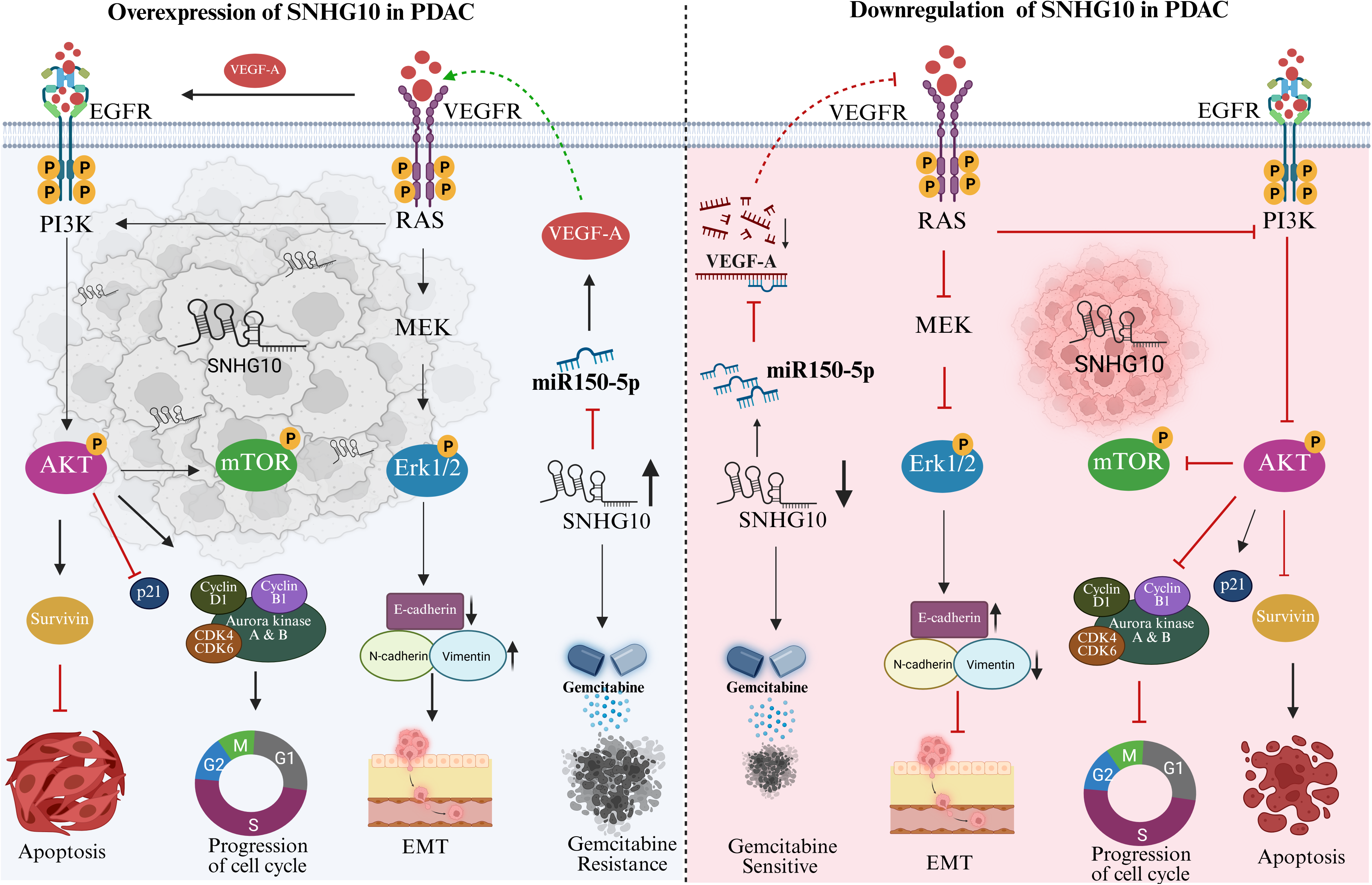

